# Recurrent connectivity supports carbon dioxide sensitivity in *Aedes aegypti* mosquitoes

**DOI:** 10.1101/2025.07.29.667487

**Authors:** Jialu Bao, Wesley Alford, Avinash Khandelwal, Laurel Walsh, George Lantz, Santiago Poncio, Laia Serratosa Capdevila, Yervand Azatian, Brian DePasquale, David G C Hildebrand, Meg A Younger, Wei-Chung Allen Lee

## Abstract

The mosquito *Aedes aegypti*’s human host-seeking behavior depends on the integration of multiple sensory cues. One of these cues, carbon dioxide (CO2), gates odorant and heat pathways and activates host-seeking behavior. The neuronal circuits underlying processing of CO2 information remain unclear. We used automated serial-section transmission electron microscopy (EM) to image and reconstruct the circuitry of the glomeruli that are innervated by the *Ae. aegypti* maxillary palp, including the glomerulus that responds to CO2. Notably, CO2-sensitive olfactory sensory neurons (OSNs) make high levels of recurrent synaptic connections with one another, while making a low density of feedforward synapses. At some of these contacts between CO2 OSNs, we observe ribbon-like presynaptic structures, which may further enhance recurrent signaling. We compared both feedforward and recurrent connectivity with all olfactory glomeruli in *Drosophila melanogaster,* and we found more recurrent connections between the *Ae. aegypti* CO2-responsive OSNs than in any *D. melanogaster* glomeruli. We developed a computational circuit model that demonstrates recurrent synapses are necessary for robust CO2 detection under normal physiological conditions. Together, elevated levels of recurrent connectivity and ribbon-like structures may amplify sensory information detected by CO2-sensitive OSNs to support mosquito activation and sensitization by CO2, even in the presence of high levels of other odorants in the environment. We propose that this circuit organization supports the salience of CO2 as a mosquito host cue.

**One Sentence Summary:** Connectomic analysis of carbon dioxide circuitry in the disease-vector mosquito *Aedes aegypti*.

## Introduction

Mosquito-borne disease claims over half a million lives every year. The mosquito *Aedes aegypti* is a globally distributed disease vector responsible for thousands of deaths annually (“Vector-borne diseases,” 2023), with fatalities projected to increase over time (Iwamura et al., 2020; Lee and Farlow, 2019). Adult female mosquitoes innately seek blood to obtain the protein required for egg maturation (Christophers, 1960) and they integrate a range of sensory cues to detect their preferred host, humans. These cues include skin-derived odorants (Dekker et al., 2005; McMeniman et al., 2014; Zhao et al., 2022), body heat (Corfas and Vosshall, 2015; Liu and Vosshall, 2019), water vapor (Khan and Maibach, 1966; Laursen et al., 2023), visual cues (Liu and Vosshall, 2019; Vinauger et al., 2019), and carbon dioxide (CO2) (Dekker et al., 2005; McMeniman et al., 2014). The host cue CO2 drives notable changes in behavior, including increased locomotion, evident from both increases in take-off and flight time, as well as increased upwind flight (Dekker et al., 2005; Dekker and Carde, 2011) These behaviors are presumed to initiate the search for a host (Dekker et al., 2005), a state change termed *activation* (Dekker et al., 2005; Dekker and Carde, 2011). An activated mosquito investigates host cues such as body odor, heat, and visual contrast. These synergize to drive host-seeking and biting behaviors, which include landing and probing with the proboscis (Dekker et al., 2005). We refer to the priming of other sensory channels during behavioral activation as CO2 *sensitization* (Dekker et al., 2005; Lacey et al., 2014) (**Figure 1A**). We sought to discover how CO2 activation and sensitization mechanisms are reflected in the neuronal circuitry that detects CO2 and integrates it with other host cues.

**Figure 1:**
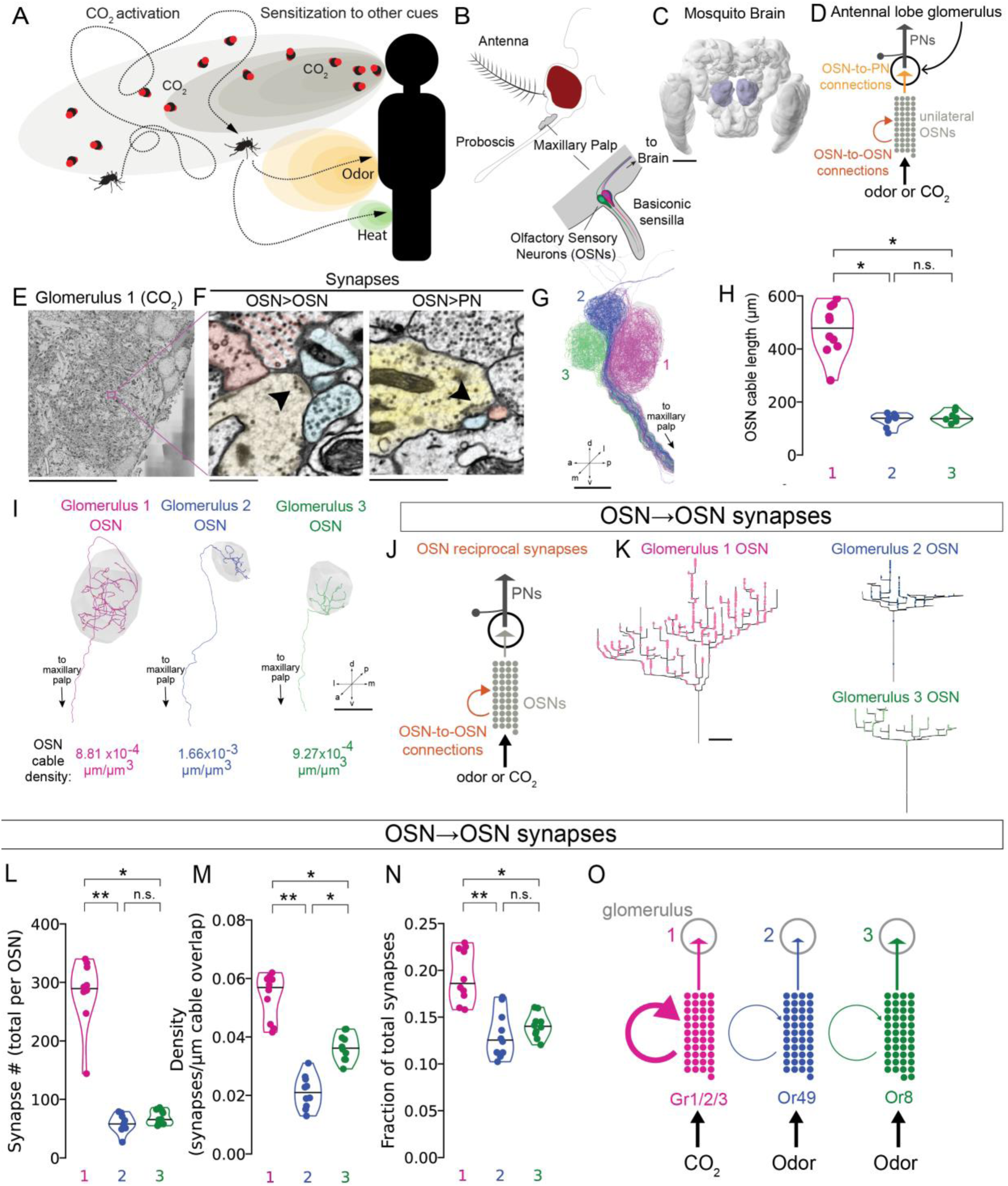
*Aedes aegypti* mosquito chemosensory driven behaviors and EM reconstruction of maxillary palp glomeruli. (A) Schematic of CO_2_ activation and sensitization in *Ae. aegypti* host-seeking. (*left*) Quiescent adult female mosquitoes become *activated* upon CO_2_ detection. (*right*) *Sensitization* to additional host cues (eg. odor, heat, etc.) drives host-seeking behavior. (B) Diagrams (*top*) of an adult female *Ae. aegypti* head and (*bottom*) a basiconic sensillum on maxillary palp (gray) which contain dendritic processes of olfactory sensory neurons (OSNs), including CO_2_-sensitive *Gr3*-expressing OSNs. OSN axons project centrally to the antennal lobes in the brain. (C) Volumetric rendering of the mosquito brain neuropils including the antennal lobes (light blue) (Heinze et al., 2021; Matthews et al., 2019). Scale bar 100 µm. (D) Schematic of an antennal lobe glomerulus and the circuit components analyzed here. Multiple OSNs selective to a chemosensory cue (eg. odor or CO_2_) converge on a glomerulus where they make synapses onto a cognate projection neuron (PN). OSNs can also make reciprocal synapses onto other OSNs. (E) Coronal 40 nm-thick section through maxillary palp Glomerulus 1, acquired and aligned using large-scale, tape-based transmission electron microscopy (EM) (Phelps et al., 2021). Scale bar 25 µm. Magenta inset expanded in (F) (*left*) Example OSN (yellow) to OSN (blue) synapse (black arrowhead). (*right*) Polyadic synapse (black arrowhead) in a Glomerulus 1 OSN (yellow). Red: postsynaptic multiglomerular cell. Blue: postsynaptic Glomerulus 1 uniglomerular projection neuron (uPN). Scale bar 1 µm. (G) Sagittal view of reconstructed maxillary palp nerve OSNs: Glomerulus 1 (pink), 2 (blue), and 3 (green). Scale bar 25 µm. a, anterior; d, dorsal; l, lateral; m, medial; p, posterior; v, ventral. (H) Cable length of OSNs (Kruskal-Wallis test (two-tailed), p = 6.20 x 10^-5^, pairwise comparisons using Dunn’s post-hoc test with Bonferroni correction: n.s., p > 0.05; * p ≤ 0.01, ** p ≤ 0.001). (I) Coronal view of individual OSNs that innervate Glomerulus 1, 2, or 3 (gray surface meshes). Scale bar 25 µm. (J) Schematic with reciprocal OSN-to-OSN connections highlighted. (K) Axograms of representative OSNs from each glomerulus and their reciprocal output synapses to other OSNs. Scale bar 25 µm. (L) Total number of outgoing OSN-to-OSN synapses contained within Glomerulus 1, 2, and 3 for ten fully reconstructed OSNs (Kruskal-Wallis test (two-tailed), p = 1.28 x 10^-5^, pairwise comparisons using Dunn’s post-hoc test with Bonferroni correction: n.s., p > 0.05, * p ≤ 0.05; ** p ≤ 0.001). (M) Synapse density of outgoing OSN-to-OSN synapses per micrometer of cable overlap (length of postsynaptic OSN axon within 2 µm of presynaptic OSN axon, see Methods) (Kruskal-Wallis test (two-tailed), p = 0.0003, pairwise comparisons using Dunn’s post-hoc test with Bonferroni correction: n.s., p > 0.05; * p ≤ 0.05; ** p ≤ 0.001). (N) Reciprocal OSN-to-OSN synapses as a fraction of overall OSN output synapses (Kruskal-Wallis test (two-tailed), p = 0.0002, pairwise comparisons using Dunn’s post-hoc test with Bonferroni correction: n.s., p > 0.05, * p ≤ 0.01, ** p ≤ 0.001). (G) Schematic of reciprocal connectivity with arrow thickness reflecting synapse number.

Many principles of chemosensory wiring and coding were established in the model fly, *Drosophila melanogaster*. Olfactory sensory neurons (OSNs) in the antennae and maxillary palps (Couto et al., 2005; Fishilevich and Vosshall, 2005) innervate ∼50 glomeruli in the antennal lobes (ALs) (Stocker et al., 1990, 1983), the first olfactory processing center of the brain. Multiple OSNs express a single olfactory receptor and all OSNs expressing the same receptor converge on a single glomerulus (Couto et al., 2005; Fishilevich and Vosshall, 2005), which can be defined as an anatomically distinct local circuit. Local neurons (LNs) project among antennal lobe glomeruli, transforming information through a combination of excitatory and inhibitory interactions (Wilson, 2013). From antennal lobe glomeruli, projection neurons (PNs) send their axons to higher brain centers. Synapse-resolution reconstructions of the *D. melanogaster* olfactory system (Gruber et al., 2023; Horne et al., 2018; Tobin et al., 2017) and connectomes of the whole brain have recently been generated (Dorkenwald et al., 2024; Scheffer et al., 2020; Schlegel et al., 2024; Zheng et al., 2018). This work revealed circuit mechanisms of sensory coding (Bates et al., 2020; Frechter et al., 2019; Schlegel et al., 2021), innate behavior (Dolan et al., 2019; Huoviala et al., 2020), and learning and memory (Dolan et al., 2019; Felsenberg et al., 2018). *Ae. aegypti* and *D. melanogaster* are both dipteran insects, but they have evolved dramatically different feeding strategies during the more than ∼150-million-year timespan since they shared a common ancestor (Arensburger et al., 2010; da Silva et al., 2020). This has led to different specializations in chemosensory information processing. Recent work identified the co-expression of multiple olfactory receptors in many *Ae. aegypti* OSNs (Fernández-Chiappe et al., 2025; Herre et al., 2022), and it is likely that additional differences in the chemosensory circuit architecture support their diet of primarily human blood.

Information processing in the olfactory system, like other neuronal circuits, relies on synapses, which are the main sites of information transmission between neurons. At chemical synapses, neurotransmitters are packaged into synaptic vesicles and released from specialized presynaptic active zones for detection by receptors at postsynaptic sites. Vesicular release machinery traffics vesicles to the presynaptic membrane, docks vesicles at the active zone, and drive vesicular exocytosis (Harris and Littleton, 2015; Südhof, 2004). The structure of presynaptic active zones is diverse across species and synapse types, and related to functional properties of different synapses (Matthews and Fuchs, 2010; Prokop and Meinertzhagen, 2006; Sterling and Matthews, 2005; Südhof, 2012). In insects, synapses typically have a small electron-dense structure referred to as a T-bar (which appears as a T when cut in cross-section) that promotes vesicle docking and release (Prokop and Meinertzhagen, 2006; Wagh et al., 2006). In contrast, vertebrate hair cells and neurons in the retina have large presynaptic specializations called ribbon synapses, which are elongated ribbon-shaped electron-dense structures extending intracellularly from the presynaptic active zone. Ribbon synapses are thought to support sustained and graded synaptic vesicle release (Matthews and Fuchs, 2010; Sterling and Matthews, 2005). Ribbon synapses include the vertebrate-specific protein Ribeye (Kantardzhieva et al., 2012; Magupalli et al., 2008; Schmitz et al., 2000) and have not previously been identified in invertebrates. The existence of ribbon-like synapses in invertebrates would imply a convergently evolved mechanism for sustained or graded synaptic vesicle release across species.

In *Ae. aegypti*, the maxillary palp (**Figure 1B**) is the sensory appendage that contains neurons that sense CO2. To examine the circuit mechanisms that drive CO2 sensitivity, we generated a synapse-resolution wiring diagram of the three glomeruli that are innervated by maxillary palp OSNs. We used automated transmission electron microscopy (EM) (Phelps et al., 2021) to generate an EM volume that includes all three glomeruli innervated by the OSNs that arise from the maxillary palp. For each of the three glomeruli, we reconstructed and annotated every synapse in 10 OSNs and their cognate uniglomerular PNs (uPNs) that exit these glomeruli and project in the inner antennocerebral tract (iACT). We found that CO2-sensitive OSNs had high levels of recurrent connections, a motif that may be used to amplify CO2 detection. To compare chemosensory coding strategies in animals with divergent feeding behaviors, we analyzed the recurrent and feedforward connectivity of all *D. melanogaster* olfactory glomeruli including the CO2 glomerulus, Glomerulus V, using FlyWire (Dorkenwald et al., 2024). We did not find recurrent or feedforward connectivity to be similar between CO2-sensitive glomeruli in *Ae. aegypti* and *D. melanogaster*. Rather, there were far more recurrent connections between CO2-sensitive OSNs in *Ae. aegypti* than the OSNs that innervate any glomerulus in the fly. We identified a novel structure located specifically in CO2-sensitive OSNs reminiscent of mammalian ribbon synapses (Kindt and Sheets, 2018; Matthews and Fuchs, 2010; Nouvian et al., 2006; Raviola and Dacheux, 1987; Sterling and Matthews, 2005), and propose that these ribbon-like structures may supplement excitatory axo-axonic signaling. Lastly, we adapted a biophysical model from *D. melanogaster* to include recurrent connections. Using this model and previously published measurements we found that recurrent connections supported robust CO2 detection, even in the presence of strong background odors that otherwise limit CO2 detection sensitivity. Together, this work suggests that elevated levels of recurrent connectivity and ribbon-like synapses may amplify incoming CO2 cues to support mosquito activation and sensitization to CO2.

## Results

### EM reconstruction of olfactory sensory neurons reveals high recurrent connectivity among CO2 neurons

*Ae. aegypti* rely on multiple sensory cues to locate humans (**Figure 1A**), and some of these cues, including CO2, are first detected in the maxillary palp (**Figure 1B**) by OSNs that innervate the antennal lobes (**Figure 1C-D**). To compare the circuits that detect and process CO2 to those dedicated to other odorants, we set out to image and reconstruct multiple antennal lobe glomeruli with electron microscopy (EM). We used automated serial-section transmission EM (Phelps et al., 2021) to image the posterior two-thirds of the antennal lobes (**Figure 1C-F**) of an adult female *Ae. aegypti* brain (**Supplementary** Figure 1A) at 4 × 4 × 40 nm^3^/voxel resolution. This region of the antennal lobe includes the three glomeruli that are innervated by the maxillary palp: the CO2-responsive glomerulus, Glomerulus 1, and two other olfactory glomeruli, Glomeruli 2 and 3, which respond to odorants (Herre et al., 2022; Ignell et al., 2005).

In order to analyze the connectivity of these circuits, we manually reconstructed neurons in Glomeruli 1, 2, and 3 on the right side of the brain (**Figure 1G-J**). We identified neurons as olfactory sensory neurons (OSNs), uniglomerular projection neurons (uPNs), or multiglomerular neurons (MGs) by their morphology, axon tracts, and which glomeruli they innervated. Most MGs extended beyond the imaged volume, and we therefore could not determine their cell type. We therefore focus our analysis on OSN and uPN circuitry (see Methods for details).

To examine the morphology of individual OSNs, we first annotated the processes of every OSN projecting from the right maxillary palp nerve by creating a skeleton reconstruction of each OSN that includes its all branches (**Figure 1G,I**). We identified 136 OSNs that innervate Glomeruli 1, 2, and 3. Each basiconic sensilla on the maxillary palp contains 3 OSNs (Mclver, 1982), and each OSN is thought to project to a single glomerulus. We found that 45 OSNs that project to Glomerulus 1, 45 to Glomerulus 2, and 46 to Glomerulus 3. Consistent with previous work (Herre et al., 2022; Ignell et al., 2005), we saw that the axonal arbors of each OSN projected to a single glomerulus (**Figure 1G,I; Supplementary** Figure 3-4).

From this population of OSNs, we chose ten OSNs innervating each glomerulus that represented the morphologies of the total population by visual inspection (**Figure 1G,I; Supplementary** Figure 3). We then tested whether these were representative of the morphologies of the total OSN population using NBLAST (Costa et al., 2016) and found that they were not significantly different from the overall population (Welch’s t-test, p= 0.21, p= 0.07, p=0.19 for Glomerulus 1, 2, and 3, respectively). The number of excitatory synapses between two cells is indicative of the amplitude and probability of depolarization in a postsynaptic neuron after the presynaptic neuron fires (Holler et al., 2021; Tobin et al., 2017). We therefore reconstructed these 30 OSNs (10 per glomerulus) to completion, meaning that all synapses made by these neurons onto any postsynaptic partner were annotated. We quantified the length of all neurites (cable length) of these fully reconstructed OSNs and found that Glomerulus 1 OSN cable length was over twice as long compared to Glomerulus 2 or 3 OSNs (**Figure 1H; Supplementary** Figure 5A). Due to the large volume of Glomerulus 1, its neurite density is lower than Glomerulus 2 or 3 despite the elevated cable length (**Figure 1H-I**).

One mechanism for signal amplification in neural networks is recurrent or reciprocal excitatory connectivity (Douglas et al., 1995; Nicoll, 1971). We next asked if *Ae. aegypti* OSNs have recurrent connectivity that could amplify the detection of host cues. We examined recurrent connections between the OSNs that innervate each glomerulus (**Figure 1J-N**). We define a recurrent synapse as any OSN-to-OSN synapse where both OSNs send axons to the same glomerulus. We found that Glomerulus 1 OSNs had 283.4 ± 54.3 recurrent synapses per neuron, while Glomerulus 2 and 3 OSNs had only 58.9± 14.5 and 68.7 ± 11.1, respectively (**Figure 1L; Supplementary** Figure 5D). We considered the possibility that there are more recurrent synapses in Glomerulus 1 OSNs simply because their processes are longer than Glomerulus 2 or 3 OSNs (**Figure 1I**). Because the opportunities for neurons to be synaptically connected are limited to locations where axons and dendrites are sufficiently close to make synapses, we computed the density of recurrent synapses per micrometer of overlapping axonal cable length (when axons are within 2 µm of each other or cable overlap) in each glomerulus (**Figure 1M**). Glomerulus 1 OSNs had the highest recurrent synapse density with 0.053 ± 0.008 synapses per micron, while Glomerulus 2 OSNs had 0.021 ± 0.006 and Glomerulus 3 OSNs had 0.036 ± 0.005 (**Figure 3D; Supplementary** Figure 5E). To account for the difference in neurite length, we also computed the fraction of the total OSN output synapses that are recurrent synapses. We also found that Glomerulus 1 OSNs had the highest fraction of recurrent synapses (**Figure 1N**). The higher number of recurrent connections among Glomerulus 1 OSNs may provide a neural substrate for amplifying excitation by CO2 cues in the mosquito brain (**Figure 10**).

### CO2-sensitive OSNs have more recurrent than feedforward synapses

To study feedforward connectivity, we reconstructed antennal lobe uPNs (**Figure 2A**). We identified the uPNs that innervate Glomerulus 1, 2, and 3, which have large-caliber axons that project dorsally in the inner antennocerebral tract (iACT). These were classified as iACT uPNs because of their axonal projections and because their dendrites did not extend beyond the glomerulus (**Supplementary** Figure 1B). We found that only one ipsilateral iACT uPN innervated each glomerulus (**Figure 2B-C**). As with the OSNs, the Glomerulus 1 uPN cable length was greater, but the neurite density was lower than for Glomerulus 2 and 3 uPNs (**Figure 2B**).

**Figure 2:**
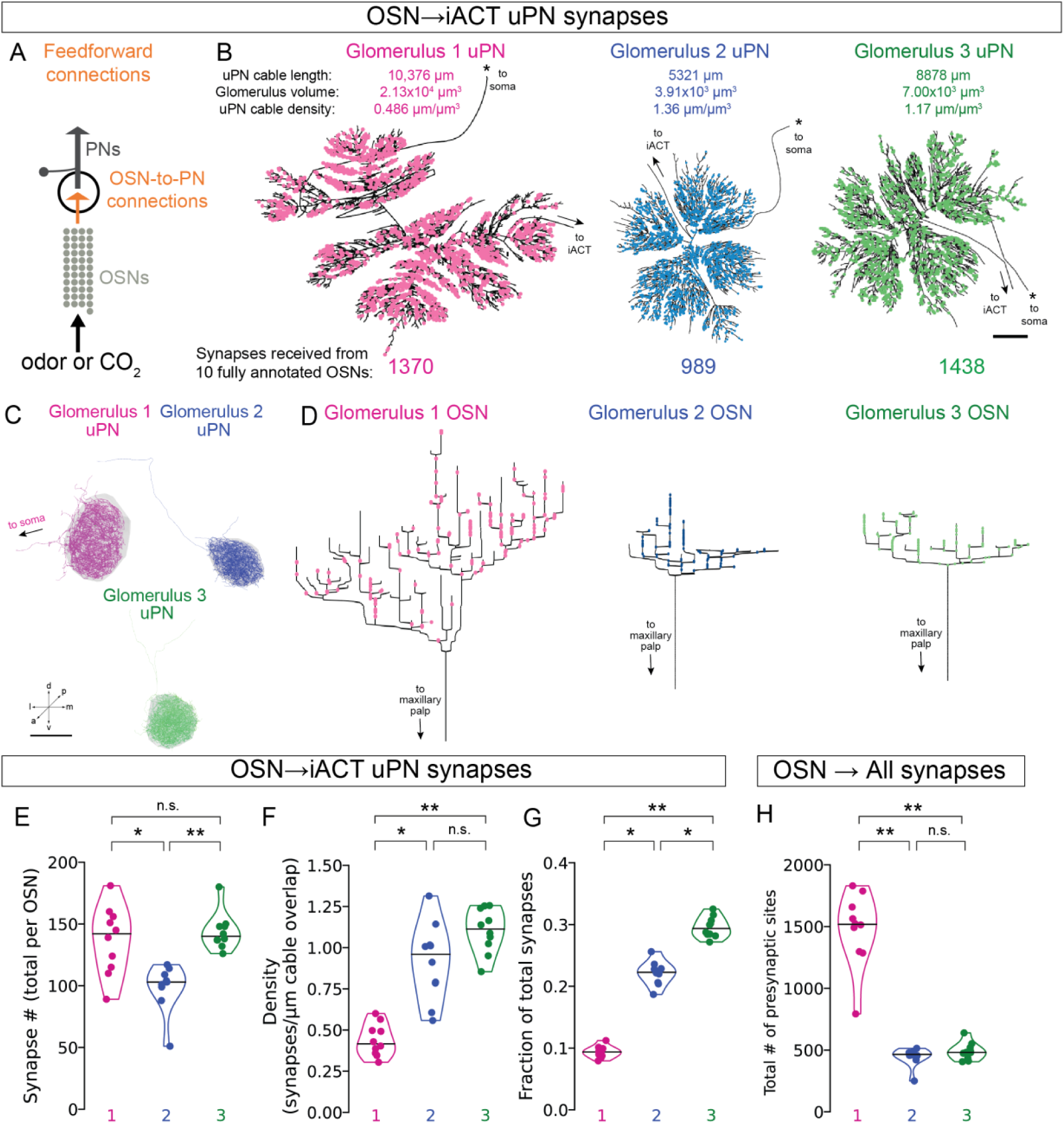
Feedforward connectivity of maxillary palp glomeruli. (A) Schematic with feedforward OSN-to-PN connections highlighted. (B) Dendrograms (flattened 2D representations of the 3D dendritic arbor preserving distance information) of uPNs and their incoming feedforward synapses from OSNs. * denotes branch leading to soma, and arrow denotes branch leading to the inner antennocerebral tract (iACT). (*top*) uPN cable length, glomerular volume, and cable density. Scale bar 100 µm. (C) Coronal view of uPN reconstructions. Scale bar 25 µm. (D) Axograms (flattened 2D representations of the 3D axonal arbor preserving distance information) of representative OSNs with their feedforward synapses to uPNs shown as colored dots. Scale bar 25 µm. (E) Number of OSN-to-uPN synapses from ten fully reconstructed OSNs within Glomerulus 1, 2, and 3 (Kruskal-Wallis test (two-tailed), p = 0.0004, pairwise comparisons using Dunn’s post-hoc test with Bonferroni correction: n.s, p > 0.05; * p ≤ 0.05; ** p ≤ 0.001). (F) Density of OSN-to-uPN synapses per µm of cable overlap (length of uPN dendrite within 2 µm of OSN axon, see Methods) (Kruskal-Wallis test (two-tailed), p = 3.34 x 10^-5^, pairwise comparisons using Dunn’s post-hoc test with Bonferroni correction: n.s., p > 0.05; * p ≤ 0.01; ** p ≤ 0.001). (G) Feedforward OSN to uPN synapses as a fraction of overall OSN output synapses (Kruskal-Wallis test (two-tailed), p = 2.49 x 10^-6^, pairwise comparisons using Dunn’s post-hoc test with Bonferroni correction: * p ≤ 0.05, ** p ≤ 0.001). (H) Total number of outgoing OSN synapses to all cell types (Kruskal-Wallis test (two-tailed), p = 4.5 x 10^-5^, pairwise comparisons using Dunn’s post-hoc test with Bonferroni correction: n.s., p > 0.05, * p ≤ 0.01; ** p ≤ 0.001).

To determine the number of feedforward synapses formed between OSNs and uPNs, we annotated all synapses onto each ipsilateral iACT uPN (**Figure 2A-C**). We found that each Glomerulus 1 OSN (**Figure 2D-E**), made 137.0 *±* 26.0 (mean *±* SD) synapses onto the Glomerulus 1 uPN. This uPN received 1370 synapses from the 10 fully annotated OSNs, and therefore receives an estimated 6000-7000 synapses from all OSNs. Although the Glomerulus 1 OSNs had larger and more complex arbors than Glomerulus 3 OSNs (**Figure 2B**), Glomerulus 1 and 3 uPNs received a comparable number of synapses from each OSN, 137.0 *±* 26.0 and 143.8 *±* 14.0 per OSN, respectively (**Figure 2E**). Glomerulus 2 uPNs received the lowest number of feedforward synapses (98.9*±* 17.7). To account for the large volume of Glomerulus 1, we again calculated the feedforward connectivity as the density of feedforward synapses per micrometer of axon-to-dendrite cable (**Figure 2F**), the fraction of the total output synapses (**Figure 2G**), and the total number of OSN output synapses (**Figure 2H**). Glomerulus 1 OSNs had the lowest density and fraction of feedforward synapses. Taken together, the Glomerulus 1 OSNs made more recurrent synapses than feedforward synapses onto the uPN that we sampled, while Glomerulus 2 and 3 OSNs made more feedforward than recurrent synapses (**Figures 1L, 2E**).

### High levels of recurrent connectivity are specific to Ae. aegypti CO2 OSNs

While CO2 initiates an activated search state in female *Ae. aegypti* mosquitoes, it has a different impact on *D. melanogaster* behavior. For *D. melanogaster*, high concentrations of CO2 are aversive when the fly is at rest, and moderate levels of CO2 can be attractive when flies are already in motion (Faucher et al., 2006; Lin et al., 2013; Suh et al., 2004; van Breugel et al., 2018; Wasserman et al., 2013). Although most of the olfactory systems of both insects utilize ORs and IRs, CO2 is detected by a gustatory receptor (GR) that is composed of Gr1, Gr2, and Gr3 subunits in *Ae. aegypti*, which are orthologues of Gr21a and Gr63a subunits in *D. melanogaster* (Jones et al., 2007; Kwon et al., 2007). We sought to determine if the organization of the CO2 glomerulus is dictated by the receptor expressed in its OSNs or if ethological significance is more predictive of circuit structure. To this end, we compared circuit features of the *Ae. aegypti* CO2 glomerulus to all *D. melanogaster* olfactory glomeruli, including the CO2 glomerulus V (**Figures 3-4**).

**Figure 3:**
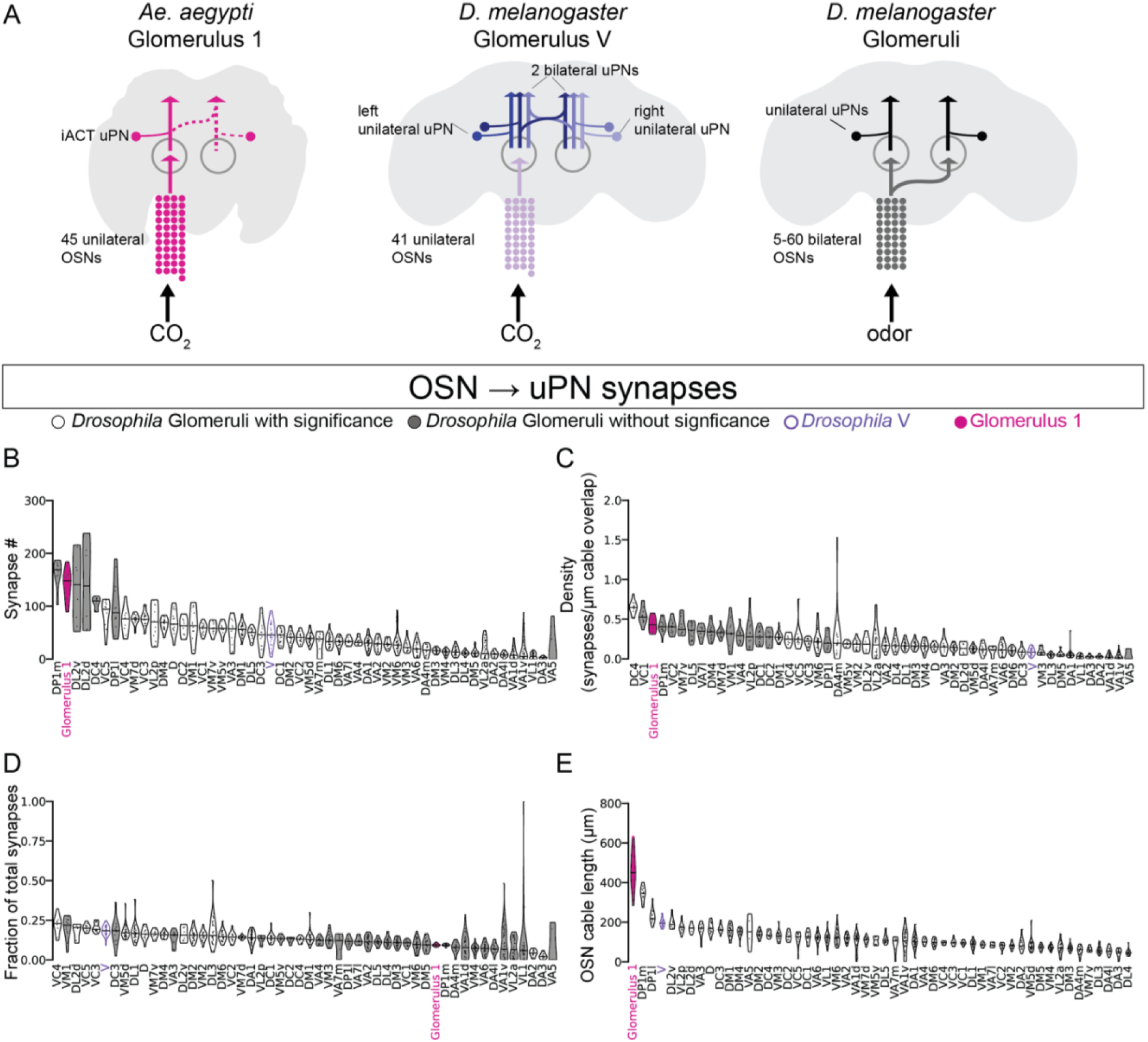
Feedforward connectivity of *Ae. aegypti* Glomerulus 1 compared to *D. melanogaster* antennal lobe glomeruli. (A) Schematics of cells and connectivity compared between *Ae. aegypti* and *D. melanogaster*. *(left)* The *Ae aegypti* CO_2_-sensitive Glomerulus 1 with 45 unilateral OSNs and the 1 uPN that projects ipsilaterally through the iACT to higher brain regions. Dashed lines indicate areas beyond the main reconstruction. *(middle)* The *D. melanogaster* CO_2_-sensitive glomerulus V with 41 unilateral OSNs and 3 uPNs: one unilateral projecting uPN and two bilateral projecting uPNs, with one bilateral uPN on each side of the brain. *(right)* Representation of *D. melanogaster* glomeruli that contain a single uPN. These contain between 5 and 60 bilateral OSNs, which each innervate the corresponding glomerulus on both sides of the brain and have 1 uPN that projects ipsilaterally to higher brain regions. (B) Number of feedforward OSN-to-uPN synapses for each glomerulus (Kruskal-Wallis test (two-tailed), p = 3.93 x 10^-56^). Filled data are non-significantly different from Glomerulus 1 using Dunn’s post-hoc test with Bonferroni correction: p ≤ 0.01). (C) Density of feedforward OSN-to-uPN synapses per µm of cable overlap within each glomerulus (Kruskal- Wallis test (two-tailed), p = 1.39 x 10^-45^. Filled data N.S. are non-significantly different using Dunn’s post-hoc test with Bonferroni correction: p ≤ 0.01). (D) Feedforward OSN-to-uPN synapses as a fraction of overall OSN output synapses (Kruskal-Wallis test (two-tailed), p = 4.87 x 10^-38^. Filled data are non-significantly different using Dunn’s post-hoc test with Bonferroni correction: p ≤ 0.01). (E) OSN axonal cable length within each glomerulus. (Kruskal-Wallis test (two-tailed), p = 1.03 x 10^-62^. Filled data are non-significantly different using Dunn’s post-hoc test with Bonferroni correction: p ≤ 0.01).

**Figure 4:**
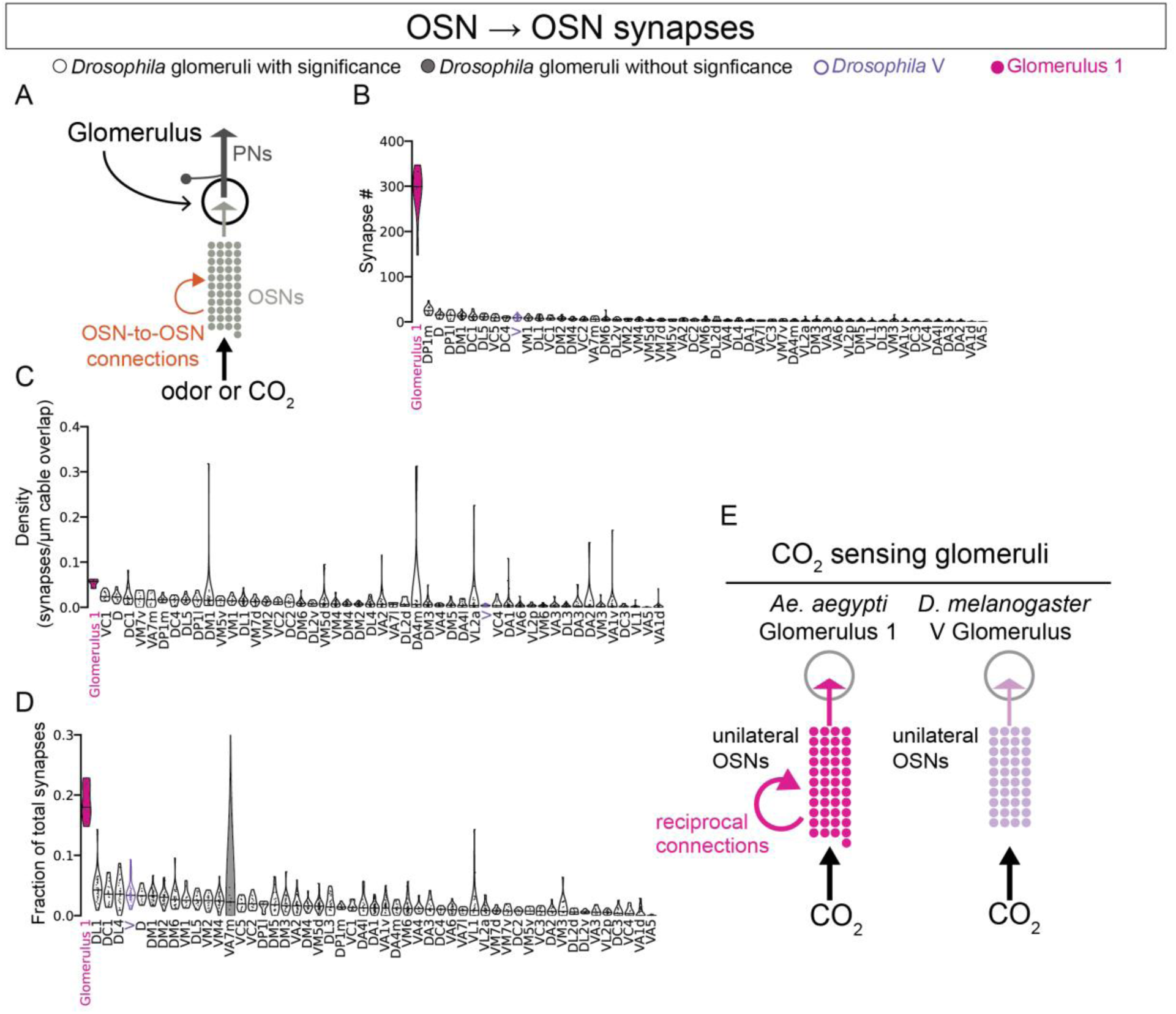
*Ae. aegypti* Glomerulus 1 has elevated reciprocal OSN-to-OSN connectivity. (A) Schematic with reciprocal OSN-to-OSN connections highlighted. (B) Total number of outgoing OSN-to-OSN synapses contained within each glomerulus (Kruskal-Wallis test (two-tailed), p = 1.63 x 10^-98^. Ten fully reconstructed OSNs are included for *Ae. aegypti* CO_2_ sensitive Glomerulus 1. Filled data are non-significantly different using Dunn’s post-hoc test with Bonferroni correction: p ≤ 0.01). (C) Density of reciprocal OSN-to-OSN synapses per µm of cable overlap within each glomerulus (Kruskal-Wallis test (two-tailed), p = 1.39 x 10^-72^. Filled data are non-significantly different using Dunn’s post-hoc test with Bonferroni correction: p ≤ 0.01) (D) Reciprocal OSN-to- OSN synapses as a fraction of overall OSN output synapses (Kruskal-Wallis test (two-tailed), p = 1.13 x 10^-72^. Filled data are non-signficantly different using Dunn’s post-hoc test with Bonferroni correction: p ≤ 0.01). (E) Schematic summarizing difference between CO_2_ selective OSN connectivity between *Ae. aegypti* and *D. melanogaster*.

We first compared feedforward synaptic connections between *Ae. aegypti* Glomerulus 1, and all *D. melanogaster* glomeruli. We leveraged our *Ae. aegypti* dataset and the *D. melanogaster* Full Adult Fly Brain dataset (Zheng et al., 2018) using segmentations and synapse predictions available in FlyWire (Dorkenwald et al., 2023, 2022). OSN projections to all *Ae. aegypti* glomeruli are unilateral, as are OSN projections to *D. melanogaster* glomerulus V. All other *D. melanogaster* olfactory glomeruli receive feedforward input from bilateral OSNs (**Figure 3A; Supplementary** Figure 6-7). To account for these morphological differences, we restricted our analysis to neurites and synapses within each right glomerulus in FlyWire. To compare feedforward connectivity across these glomeruli, we quantified the number of synapses each OSN makes onto its cognate uPNs (**Figure 3B**). We found that the feedforward synapse number (**Figure 3B**), feedforward synapse density (**Figure 3C**). and feedforward fraction of synapses (**Figure 3D**) of Glomerulus 1 were similar to many of the fly glomeruli, despite the large size of this glomerulus, however, the neurite length was greater than any of the fly glomeruli (**Figure 3E**). The total number of feedforward synapses in Glomerulus 1 was on the high end of the distribution, but was not significantly different from some of the fly glomeruli, including DC4 and DP1m, which have both been implicated in acid sensing (Ai et al., 2013, 2010).

The *D. melanogaster* V glomerulus is innervated by dendrites from a unilateral uPN in each hemisphere, and two bilateral uPNs, one with a soma in the left hemisphere and one in the right hemisphere (**Figure 3A**). In order to compare feedforward OSN-to-uPN connectivity between Glomerulus 1 and the fly V glomerulus, we included the three uPNs that innervate the right V glomerulus in the analysis: the right uPN, right bilateral uPN, and left bilateral uPN (**Supplementary** Figure 9). In glomerulus V, the unilateral uPN received 5.4 ± 4.0 synapses from each OSN, and the right and left bilateral uPNs received 22.4 ± 10.0 and 19.9 ± 10.3, respectively (**Supplementary** Figure 9). The total number of feedforward synapses was 47.7 ± 22.1, which was lower in number (**Figure 3B**) and density (**Figure 3C**) than Glomerulus 1. The total fraction of feedforward synapses in V was higher than in Glomerulus 1 (**Figure 3D)**. In summary, although Glomerulus 1 and V respond to the same sensory cue, the feedforward connectivity of Glomerulus 1 does not resemble V relative to other fly olfactory glomeruli.

We next compared recurrent OSN connectivity in *D. melanogaster* glomeruli to Glomerulus 1 (**Figure 4A**). We found that Glomerulus 1 OSNs made ∼11-times more recurrent synapses (283.4 ± 53.4) than OSNs of any fly glomeruli (**Figure 4B**), and Glomerulus 1 had elevated OSN-to-OSN synapse density (**Figure 4C**) and fraction (**Figure 4D**) compared to *D. melanogaster* glomeruli. V was not similar to Glomerulus 1 in any of these metrics. We also found that all three *Ae. aegypti* maxillary palp glomeruli had higher numbers of recurrent connections than the fly glomeruli (**Supplementary** Figure 8), although we note that none compared to the high levels seen in Glomerulus 1. Taken together, Glomerulus 1 has comparable levels of feedforward connectivity, but far more recurrent connectivity than olfactory glomeruli in *D. melanogaster,* including the CO2-sensing V glomerulus (**Figure 4E**).

### Putative ribbon-like synapses are specifically between Ae. aegypti CO2 sensory neurons

Synapses in *D. melanogaster*, and most other insects are typified by presynaptic structures called T- bars, named for their recognizable “T” shaped presynaptic density when viewed in cross section (Prokop and Meinertzhagen, 2006; Wagh et al., 2006). We were surprised to identify two morphologically distinct categories of electron-dense specializations in *Ae. aegypti*. The majority of presynaptic structures resembled T-bars in the fruit fly (**Figure 5A,D**), although in *Ae. aegypti* the base or “pedestal” of the T-bars appear shorter in our specimen. The “table-tops” of the T-bars are oriented parallel to the presynaptic membrane and are lined with synaptic vesicles on the side distal to the membrane. T-bars make up the vast majority of synapses in our dataset and occur between all combinations of cell types we examined.

**Figure 5:**
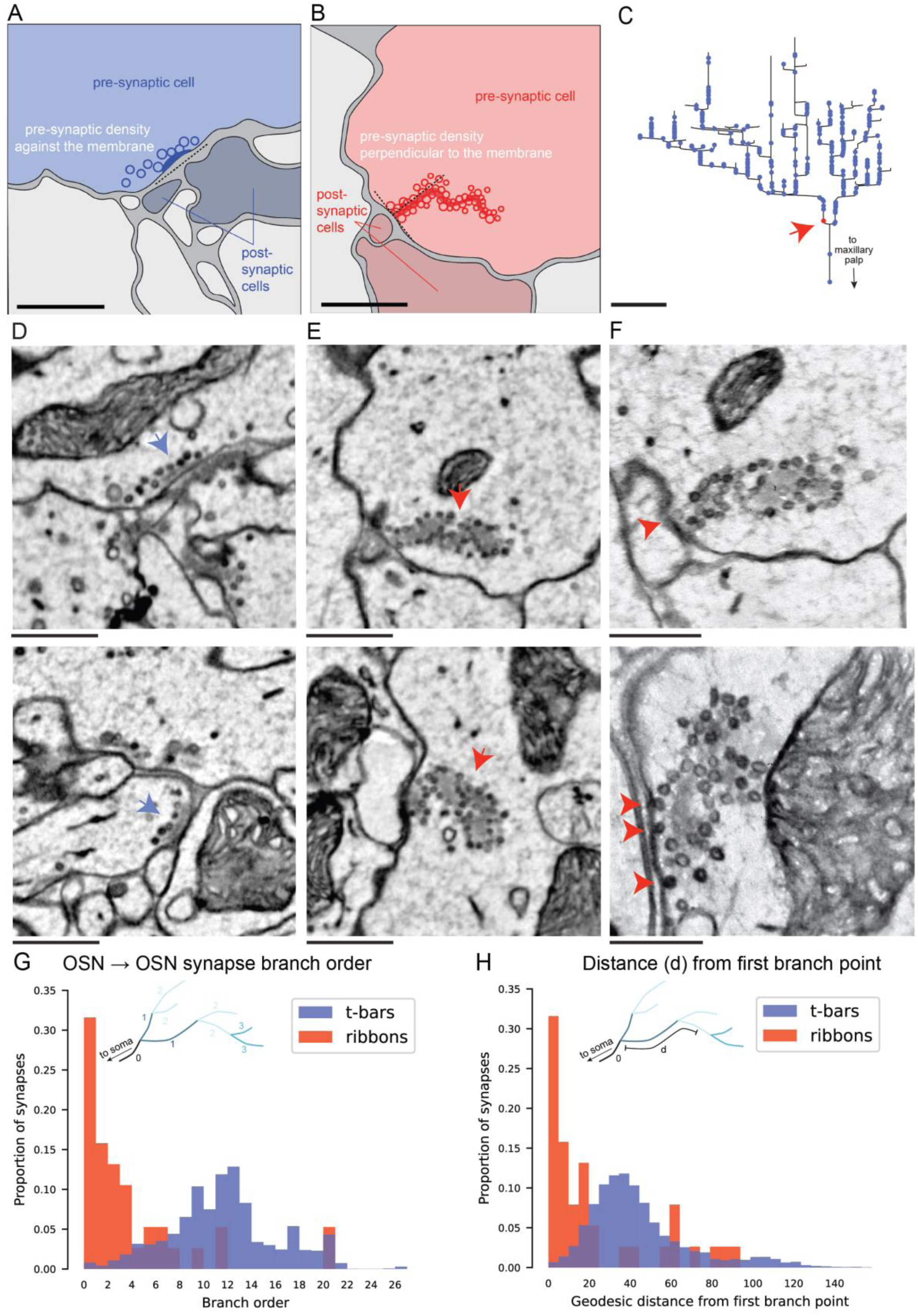
Ribbon-like synapses among CO_2_ OSNs. (A) Schematic of a canonical T-bar synapse and (B) a putative ribbon-like synapse in *Ae. aegypti* Glomerulus 1. Scale bars 500 nm. (C) Axogram of an example Glomerulus 1 OSN with reciprocal output T-bar (blue) and putative ribbon-like (red) synapses represented by colored dots. Arrow points to a putative ribbon-like synapse from this OSN. Scale bar 25 µm. (D) EM micrographs of canonical T-bar (blue arrows) synapses. Scale bars 500 nm. (E) EM micrographs of putative ribbon-like (red arrows) synapses. Scale bars 500 nm. (F) Higher magnification (0.5 nm/pixel) EM micrographs of putative ribbon-like synapses with vesicles docked to the presynaptic membrane (arrowheads). Scale bars 200 nm. (G) Histogram of putative ribbon-like and T-bar OSN-to-OSN synapse branch order (Mann-Whitney U test, p = 5.93 x 10^-17^). Inset: illustration of synapse branch order. (H) Histogram of distances (cable length) from the first branch point to putative ribbon-like and T-bar OSN-to-OSN synapses (Mann-Whitney U test, p = 6.78 x 10^-9^). Inset: illustration of quantified distance (geodesic distance or cable length) of synapse from the first branch point.

We also found a number of electron-dense structures that were oriented perpendicularly to the cell membrane and resemble ribbon synapses found in mammalian retina and vertebrate hair cells, which are named for their elongated ribbon-like morphology (**Figure 5B,E; Supplementary** Figures 10-11) (Kindt and Sheets, 2018; Matthews and Fuchs, 2010; Nouvian et al., 2006; Raviola and Dacheux, 1987; Sterling and Matthews, 2005). These densities were lined with vesicles on both sides. These putative ribbon-like synapses were only seen in OSNs that innervate Glomerulus 1 and are only located at recurrent contacts between CO2-sensitive OSNs. Within the reconstructed OSNs we saw a total of 60 ribbon-like specializations, around 30 on each side of the brain. These ribbon-like structures are located most often near the primary branches of OSNs close to the point where the axon first enters the glomerulus, while T-bars are found throughout the entire OSN arbor in the glomerulus (**Figure 5C,G,H**). We did not detect ribbon-like structures in OSNs or PNs in any other glomeruli we analyzed, in neither *Ae. aegypti* nor *D. melanogaster*, suggesting they may be specific to CO2-sensitive Glomerulus 1 OSNs in the mosquito.

To determine whether these ribbon-like structures could be presynaptic specializations that participate in synaptic transmission, we looked for the presence of vesicles that were “docked” at the presynaptic membrane. Docked vesicles contact the presynaptic membrane in preparation for rapid exocytosis to release neurotransmitter into the synaptic cleft (Südhof, 2004). To identify putative docked vesicles, we re-imaged 19 ribbon-like structures at higher magnification (0.32 × 0.32 nm^2^ per pixel and 0.4 × 0.4 nm^2^ per pixel). We identified instances where vesicles were in contact with the presynaptic plasma membrane (**Figure 5F**), which is a signature of docked vesicles. This supports the hypothesis that these ribbon-like structures may function as presynaptic specializations involved in synaptic transmission.

### Modeling reveals a role for recurrent connections in enhancing CO2 detection

To better understand the impact of recurrent connections on olfactory encoding, we constructed a mathematical model that captured key elements of olfactory processing (**Figure 6A**). We envisioned an otherwise standard *D. melanogaster* olfactory system, with a single specialized CO2 glomerulus with recurrent connections. Following existing approaches (Luo et al., 2010; Olsen et al., 2010) we model the OSN to PN circuit including inhibitory gain normalization. In *D. melanogaster*, inhibitory local neurons (LNs) in the antennal lobe mediate interglomerular inhibition upon odorant activation (**Figure 6A**). We considered how LN inhibition impacts Glomerulus 1 PN activity when other glomeruli, which lack recurrent connections, are activated by background odorants, thereby increasing LN activity. This model takes the following form:

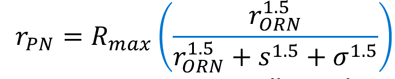

**Figure 6:**
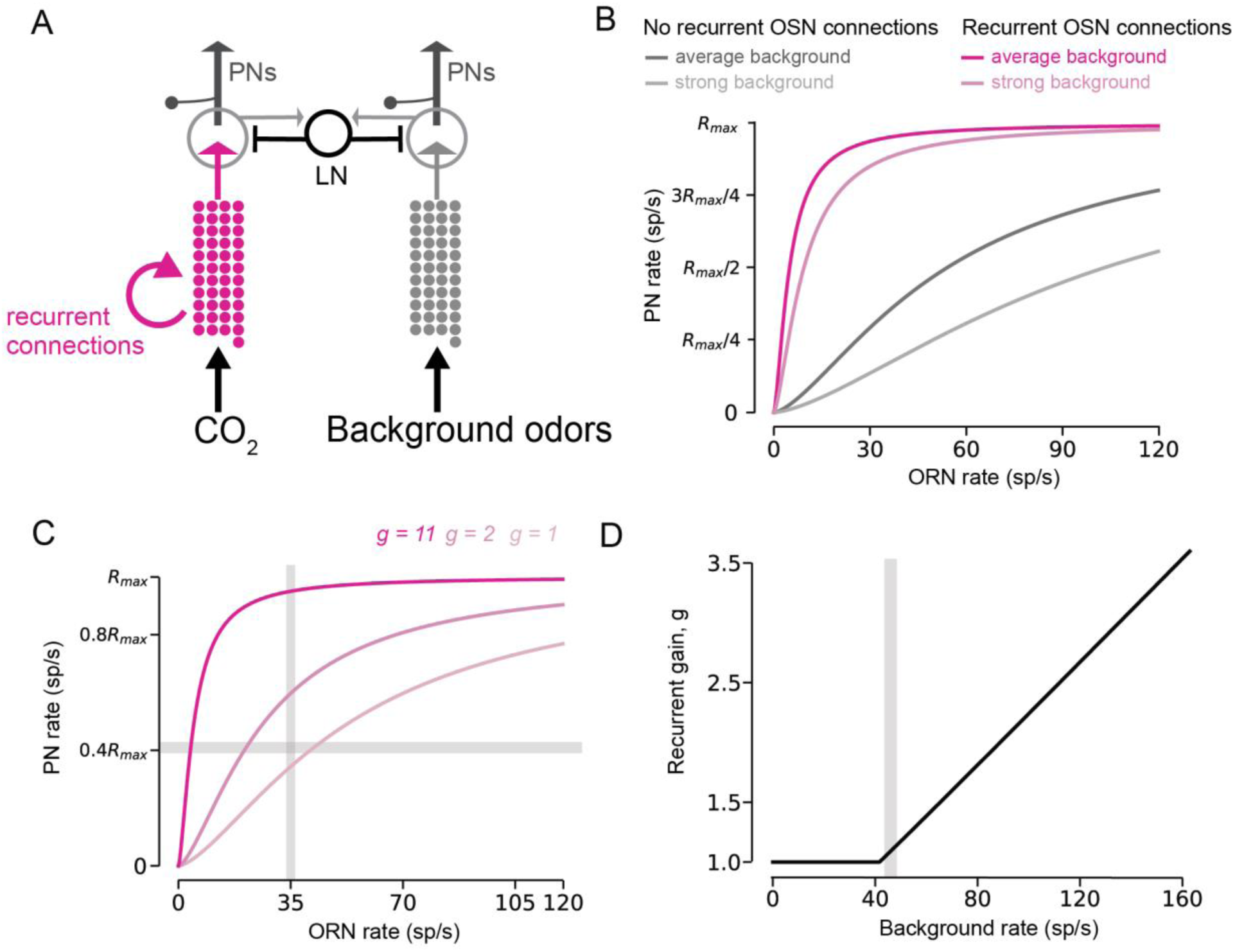
Connectomically-informed network model reveals recurrent connectivity specifically enhances stimulus detection. (A) Model architecture. CO_2_ is processed by Glomerulus 1, which includes recurrent connections. Co-occurring odorants (‘background’) activate glomeruli which inhibit other glomeruli through inhibitory local neurons (LNs). (B) OSN to PN response curves for OSNs with and without recurrent connections, for two different background odor strengths (average and strong, see Methods). (C) OSN to PN response curve for differing values of *g* for an average background odor strength. Horizontal gray shaded region indicates OSN response rates for which CO_2_ should be detected (35 sp/s), and vertical gray shaded region indicates the PN response strength that would correspond to CO_2_ detection (0.4*R_max_*). (D) Minimum value of *g* required to ensure CO_2_ detection for varying background odor strengths. Vertical gray shaded region indicates an average level of background odor.

where *Rmax* is the maximum PN response rate, *s* mediates inter-glomeruli divisive inhibition via a weighted average of all glomeruli to given odorant, and σ defines the OSN value that achieves half-max activity. Values of α, *Rmax*, and the exponent of 1.5 were adopted from (Olsen et al., 2010). Recurrent connections were incorporated as a multiplicative gain *g* applied to OSN activity (𝑟_&’"_→ 𝑔 · 𝑟_&’"_). We viewed this as the collective gain increase due to anatomical connections as well as their effective strength, which we estimate from EM data. We simulated our model under different scenarios to illustrate the impact of recurrent OSN connections on PN activity. To simulate CO2 in the presence of other odors (‘background odors’), we set *s* = 50 spikes/s, consistent with previous models (Hallem and Carlson, 2006; Luo et al., 2010; Olsen et al., 2010). We refer to background odors of this strength as ‘average’ background.

We simulated PN responses to different OSN levels when recurrent connections were present (**Figure 6B**, magenta) and were not present (**Figure 6B**, gray). Recurrent connections were simulated by increasing *g* by a factor of 11, simulating the presence of 11-times the number of recurrent connections. Recurrent connections lead to a sharp increase in the PN response curve, yielding a significantly larger PN response to smaller OSN activity. To examine the sensitivity of this gain to increases in the background odor, we doubled *s* to 100 spikes/s (**Figure 6B**, ‘strong background’), demonstrating that global OSN activity decreases individual PN response due to lateral inhibition. These results are consistent with a hypothesis that recurrent connections amplify weak OSN input for the important ethological cue of CO2.

Our primary hypothesis is that recurrent connections in Glomerulus 1 facilitate CO2 detection. To illustrate this we considered a detection task where a PN response greater than a threshold *Rthresh* = 0.4*Rmax* indicated the presence of CO2 (**Figure 6C**). For accurate CO2 detection, PN activity should reach *Rthresh* for CO2 concentrations found to elicit weak OSN responses. Based on published results (Majeed et al., 2014), we identified 35 sp/s as the OSN threshold for CO2 response detection, for which PNs should become robustly active.

We explored values of *g* that achieved a PN response greater than or equal to *Rthresh* for OSN responses equal to 35 sp/s, for an average background odor (**Figure 6C**; *s =* 50). We also identified the minimum value of *g* that achieves this for a range of background odor strengths (**Figure 6D**). For average background odor strength, both *g=11* and *g=2* (**Figure 6C**, dark and medium magenta, respectively) produce responses greater than 0.4*Rmax* for OSN values of 35 sp/s, while *g=1* (**Figure 6C**, light magenta) did not. We found that for average or stronger background odor strengths (*s=50* or greater) *g* would need to be greater than one to achieve detection of transient CO2 increases (**Figure 6D**), indicating that recurrent connections are necessary even under normal environmental conditions to ensure robust detection. Empirically, we measured a relatively large value for *g* that may suggest: (1) an extremely strong preference for detecting low levels of CO2, even at the expense of false alarms; (2) inhibition is significantly stronger than excitation in this circuit, and/or (3) the recurrent synapses we identified are effectively weak despite their number. Taken together, our modeling results demonstrate how recurrent connections between OSNs can amplify weak sensory responses to achieve robust stimulus detection even in the presence of realistic background odor levels.

## Discussion

To investigate the organizational principles of neuronal networks supporting innate sensory-driven behavior, we present the first microcircuit connectome in the *Ae. aegypti* mosquito. We used automated tape-based transmission EM (Phelps et al., 2021) and manual reconstruction to trace and annotate the wiring and connectivity of all three sensory glomeruli innervated by the maxillary palp, including the CO2-sensitive glomerulus (Glomerulus 1). We found that each glomerulus had an array of 45-46 OSN inputs, and that the OSNs that innervate the CO2-responsive Glomerulus 1 had a uniquely high level of recurrent OSN-to-OSN connectivity. Among the recurrent connections were structures that resemble ribbon synapses, which may couple OSNs to one another. We identified a single uPN that projects through the ipsilateral iACT in each palp glomerulus and analyzed feedforward connectivity onto these neurons. Using comparative connectomics, we found that the feedforward wiring of the *Ae. aegypti* CO2-sensitive glomerulus is similar to glomeruli in *D. melanogaster*, but with far higher recurrent connectivity. We found no obvious similarities between Glomerulus 1 and the V glomerulus, which share related receptors for CO2. Our results suggest that highly recurrent excitatory connectivity between CO2-sensitive OSNs in *Ae. aegypti* may amplify CO2 detection to enable behavioral activation and sensitization. We further suggest that similar features of behavioral salience may predict the architecture of microcircuits across species. In support of our experimental findings, we demonstrated with network modeling how recurrent connections improve stimulus detection under normal environmental conditions and in the face of strong background odors which might make detection harder.

### Recurrent excitatory connectivity

What is the functional role of recurrent synapses between OSNs? We predict that excitatory recurrent connections between Glomerulus 1 OSNs could support signal amplification, propagation, persistent activity, or a combination of these processes. Recurrent amplification has been reported in multiple sensory systems. In mammalian sensory cortex, excitatory pyramidal neuron responses are thought to arise from selective amplification of thalamocortical signals (Harris and Mrsic-Flogel, 2013; Li et al., 2013; Lien and Scanziani, 2013) through recurrent connections between similarly selective excitatory neurons (Cossell et al., 2015; Douglas et al., 1995; Lee et al., 2016; Oldenburg et al., 2024; Peron et al., 2020). In *D. melanogaster* a recurrent connectivity motif supports sustained activity for olfactory learning in the mushroom body (Cognigni et al., 2018). It may be surprising that amplification would occur so early in sensory processing; however, CO2 is a particularly important cue for *Ae. aegypti*, and the location of these recurrent connections would provide a mechanism to amplify CO2 signaling prior to inhibition by LNs. This may create a mechanism that could surmount the strong and broad inhibitory tone provided by LNs in the presence of other chemosensory input (Wilson and Laurent, 2005).

Reliable signal propagation across long distances is a general challenge of nervous systems (Debanne, 2004). Notably, the reconstructed OSNs and their recurrent connections are far from their sensory endings in the maxillary palp. Given that these unmyelinated axons travel a long distance along the maxillary palp nerve and subsequently branch and ramify as they enter the glomerulus, one possibility is that recurrent synapses between OSNs may provide a mechanism to mitigate signal propagation failures and increase the robustness of CO2 detection. Consistent with this idea, recurrent connections in mammalian cortex enhance the reliability of sensory signals for subnetworks with more recurrent connectivity, and loss of a few neurons in the recurrent network results in degraded stimulus encoding (Peron et al., 2020).

Recurrent connectivity has been widely associated with sustained and persistent activity, a hallmark of short-term working memory (Goldman-Rakic, 1995; Wang, 2001). In the mammalian cortex, recurrent connections are hypothesized to facilitate this process, providing a neural substrate for maintaining information over short timescales. However, examples of recurrence extend beyond the cortex. In invertebrates, one notable case is the sustained bump of activity in the ring attractor networks of the *D. melanogaster* central complex, which plays a key role in spatial navigation (Hulse et al., 2021; Kim et al., 2017). Persistent activity in the CO2 sensory system is an intriguing possibility in female *Ae. aegypti,* where CO2 elicits a prolonged activated behavioral state. CO2 sensitizes the organism and, when integrated with other host cues, drives innate host-seeking behavior that typically lasts minutes (Sorrells et al., 2022). Notably, Glomerulus 3 OSNs respond to the host odorant 1- octen-3-ol also have recurrent OSN-to-OSN connections, although fewer than between CO2-sensitive OSNs, while Glomerulus 2 OSNs have relatively few recurrent connections and are not clearly characterized as host-detecting neurons (Herre et al., 2022). The role of recurrent connections in host detection and neuron physiology remains to be seen, as virtually all mosquito OSN electrophysiology has been performed at the sensory periphery, far from the recurrent connections we describe (Kellogg, 1970).

A recent analysis of a *D. melanogaster* hemibrain connectome (Scheffer et al., 2020) reported the presence of recurrent axo-axonic connections between OSNs in *D. melanogaster* (Manoim- Wolkovitz et al., 2025). Analysis of this connectome diverged from our findings using the FlyWire connectome. In this work, network modeling demonstrated how activity-dependent recurrent connections for every glomerulus could lead to greater odor discrimination by pattern decorrelation, effectively supporting the functional role long established for divisive inhibition (Luo et al., 2010; Olsen et al., 2010). Our work demonstrates an alternative, but complementary role for recurrent connections for a single, high-priority glomerulus: odor detection via signal amplification. Further experimental investigation is necessary to understand how species-specific olfactory systems balance these two computations, perhaps as a function of ethological niche.

### Invertebrate ribbon-like synapses

We identified structures specifically between CO2-sensitive OSNs that resemble ribbon synapses seen in vertebrates. Examples of vertebrate ribbon synapses include retinal photoreceptors and bipolar cells (Raviola and Dacheux, 1987; Raviola and Raviola, 1967), auditory hair cells (Nouvian et al., 2006), and fish lateral line hair cells (Kindt and Sheets, 2018), where ribbon synapses scaffold vesicle pools that enable sustained, graded release (Matthews and Fuchs, 2010; Sterling and Matthews, 2005) (**Supplementary** Figure 10). Presynaptic ribbon-like structures have not previously been reported in insects and have not been reported in recently generated *D. melanogaster* connectomes. If these structures are indeed ribbon synapses, they would be the first identified in invertebrates, although we note that their function in neurotransmission remains to be studied.

Vertebrate ribbon synapses contain a protein encoded by the *ribeye* gene (Schmitz et al., 2000), which makes up the electron-dense structure of the ribbon (Maxeiner et al., 2016; Schmitz et al., 2000; tom Dieck et al., 2005). The Ribeye protein contains two domains: the A domain that is believed to confer ribbon structure at active zones, and the B domain that is orthologous to the transcriptional repressor with enzymatic activity, CtBP2 (Schmitz et al., 2000; Sterling and Matthews, 2005). Although there are invertebrate gene orthologues of the *CtBP2* B domain, there are no orthologues of the *ribeye* A domain that is required for ribbon assembly. The absence of the *ribeye* domain from invertebrate genomes has led to the assertion that ribbon synapses are a vertebrate specialization. We speculate that the development of ribbon-like synapses in vertebrates and invertebrates may be a case of convergent evolution. Alternatively, invertebrates may have a Ribeye- like scaffold protein encoded by a divergent DNA sequence that hinders detection with DNA alignment algorithms, but nonetheless has a similar protein structure. Further study is needed to investigate the proteins that compose these structures.

We observe ribbon-like structures exclusively at recurrent contacts between CO2-sensitive OSNs and only at the most proximal axon branches, which correspond to the entry point of axons into Glomerulus 1. The locations of these putative ribbon-like synapses suggests that they may amplify recurrent connectivity specifically among CO2 sensory neurons.

Finally, it remains possible that these are not ribbon synapses, but rather a structure that appears similar when imaged with transmission EM. It is a challenge to explore their function because they are located deep in the antennal lobe, they are in small axo-axonic connections, and we currently lack an identified protein component such as Ribeye to gain genetic access to these structures. Given their close proximity with synaptic vesicles, one alternative is that they are specializations involved in trafficking or otherwise organizing vesicles in CO2-selective OSNs. It is notable that these specializations have, to our knowledge, not been previously reported in insects and appear specifically in *Ae. aegypti* OSNs selective for CO2. Future work is needed to uncover their functional role in mosquito CO2 processing.

### Circuit logic is not based on receptor alone

*Ae. aegypti* females are obligate blood drinkers, while *D. melanogaster* are dietary generalists that feed on decaying plant matter. The two Dipteran species diverged over 150 million years ago (Arensburger et al., 2010; da Silva et al., 2020) and retain the same gross brain morphology (Ignell et al., 2005; Rajashekhar and Singh, 1994). Here, we compared the connectivity of the olfactory system of these divergent feeders at synaptic resolution. We observed a number of similarities between both organisms, including the prevalence of T-bar-like synapses as the predominant presynaptic specialization, although we note slight differences in the morphology of the T-bars. Some additional differences were clear at the onset of this study, *Ae. aegypti* OSNs send unilateral projections to the ipsilateral glomerulus while those in *D. melanogaster* largely send bilateral projections to mirror symmetric glomeruli in both the ipsi- and contra-lateral antennal lobes. One exception to this rule is the CO2-sensitive OSN projections to the *D. melanogaster* V glomerulus, which are exclusively ipsilateral. It is notable that CO2-sensitive OSNs in *Ae. aegypti* are housed in the maxillary palp, a different sensory organ than their location in the *D. melanogaster* antenna.

The role of CO2 differs between these species. In *D. melanogaster*, behavioral response to CO2 is context-dependent and can be either aversive or attractive (Faucher et al., 2006; Lin et al., 2013; Suh et al., 2004; van Breugel et al., 2018; Wasserman et al., 2013). Flies avoid regions of high CO2 concentration, which is a signature of inhospitable environments (Lin et al., 2013). CO2 is also a component of an odorant mixture that *D. melanogaster* emits when stressed, and conspecifics innately avoid this mixture (Suh et al., 2004). However, CO2 can also be attractive, as decaying food sources release this cue (Faucher et al., 2006; van Breugel et al., 2018). While *D. melanogaster* is attracted to CO2 during active foraging, attraction relies on a separate receptor and circuit that utilizes the subunit Ir25a (van Breugel et al., 2018). The feedforward connectivity of CO2 neurons in *Ae. aegypti* Glomerulus 1 and *D. melanogaster* glomerulus V at the first synaptic relay in the brain were not particularly similar compared to all other glomeruli in *D. melanogaster*. An underlying relationship between this sensory architecture and the valence or salience of sensory cues may exist, but we did not see a clear correlation. However, the recurrent connectivity of CO2-sensitive OSNs was strikingly different, suggesting differential signal processing between species.

### Beyond CO2 sensitization: host cue integration in the Ae. aegypti brain

Effective host-seeking behavior requires integration of multiple sensory cues (Corfas and Vosshall, 2015; Dekker et al., 2005; Herre et al., 2022; Laursen et al., 2023; Liu and Vosshall, 2019; McMeniman et al., 2014; Vinauger et al., 2019; Zhao et al., 2022). Our analysis of the CO2 system lays the groundwork for studying the integration of CO2 with other host cues in the *Ae. aegypti* brain. The boundaries of the EM volume analyzed here did not allow for the analysis of the complete CO2 circuit. There are likely additional uPNs and MGs that innervate these glomeruli, including a network of LNs. Our feedforward analysis is restricted to only part of the circuit, and future work to describe the entire CO2 pathway will be a major step towards understating how multiple cues are integrated in the mosquito brain. In *D. melanogaster*, PNs mainly project to two higher-order brain areas: the mushroom body, crucial for learning and memory, and the lateral horn, thought to be critical for sensory-driven innate behavior (Connolly et al., 1996; de Belle and Heisenberg, 1994; Masse et al., 2009). As Glomerulus 1 is innervated by an ipsilateral iACT-projecting uPN, we predict that multiple sensory pathways converge in the mosquito lateral horn to support innate host-seeking behavior. Examining such hypotheses awaits an *Ae. aegypti* whole-brain connectome.

## Methods

### Experimental model and subject details

*Aedes aegypti* wild-type laboratory Liverpool mosquitoes (LVPib12 strain) were maintained and reared at 25-28°C, 70-80% relative humidity, with a photoperiod of 14 hr light: 10 hr dark as previously described (Matthews et al., 2016). Adult mosquitoes were provided constant access to 10% sucrose. Male and female mosquitoes were housed together. The specimen used for this EM dataset was an adult female that was dissected nine days post-eclosion.

### Specimen preparation

Procedures involving animals were conducted in compliance with NIH guidelines and approved by the IACUC at Rockefeller University and Harvard Medical School. We fixed and stained the brain of one adult female *Aedes aegypti*. After the mosquito was immobilized on ice, the head was removed and fixed for three hours in 2.5% paraformaldehyde/2.5% glutaraldehyde in cacodylate buffer (0.1M cacodylate buffer at pH 7.4 with 0.04% CaCl2). The brain was dissected out of the head capsule in ice-cold phosphate buffer saline (PBS) and stored in fresh fixative overnight.

The dissected brain was then washed three times for ten minutes each with 0.02M 3-amino- 1,2,4-triazole (A-TRA) (van Emburg and de Bruijn, 1984; Zheng et al., 2018) in cacodylate buffer, stained in 1% osmium tetroxide, 0.1M A-TRA in cacodylate buffer for 90 minutes on ice, washed in cold cacodylate buffer, and brought up to room temperature. Following a second round of staining in 2% OsO4 (aqueous), the specimen was rinsed in filtered double distilled water (ddH2O), placed in 1% thiocarbohydrazide (aqueous) at 40°C for 8 minutes, then rinsed in ddH2O again. The brain was rinsed in maleate buffer (pH 5.15) and stained in 1% uranyl acetate in maleate buffer overnight at 4°C.

The following day, the brain was washed in maleate buffer (pH 5.15) followed by ddH2O. It was stained in lead aspartate for 3 hours at 60°C and washed in ddH2O. The brain was dehydrated in a graded ethanol series on ice, infiltrated, and embedded in resin (LX-112, Ladd Research). The embedded brain was polymerized at 60°C for ∼36 hours.

### Automated ultramicrotomy

The embedded specimen was automatically serially sectioned onto a tape-based collection substrate as described previously (Phelps et al., 2021). Briefly, the resin block with the brain was trimmed (Trim 90 knife, Diatome) into an oblong hexagonal shape. Sides of the hexagon were approximately 1000 µm, 3000 µm, and 4000 µm. The block was sectioned onto a Kapton substrate, GridTape (Luxel Corp), along the anterior-posterior axis using a customized an automated tape-collecting microtome (ATUM) (Hayworth et al., 2014; Phelps et al., 2021) system attached to an ultramicrotome (Leica UC7). We targeted 40 nm section thickness. The automatically collected sections were accurately positioned (within 200 µm from slot to slot) onto GridTape slots.

### Annotation of regions of interest

We used a custom-made tape-handling machine with in-line illumination and a reel-to-reel system to capture overview images of slots and annotate regions of interest (ROIs) containing the brain. We specified regions of interest for EM imaging using a custom MATLAB script.

### Automated transmission EM imaging

The GridTape reel containing the serial sections was placed into a custom reel-to-reel sample stage for continuous automatic acquisition (Phelps et al., 2021) on a transmission EM (JEOL 1200 EX) with a 2 × 2 array of sCMOS cameras (Zyla 4.2, Andor). Magnification was 2500× on the microscope, accelerating potential was 120 kV, and the beam current was ∼90 microamperes through a tungsten filament. Using the annotated ROIs, micrographs of 1747 sections spanning the posterior antennal lobes and including the three palp glomeruli were acquired at a resolution of 4 × 4 nm per pixel.

Across the series of 1747 imaged sections comprising the dataset, there were 29 single-section losses, with no consecutive section losses. Staining artifacts were usually small and did not adversely affect annotation, especially in large neuronal processes. Small, <1 µm membrane breaks occurred in some large caliber axonal processes, but were straightforward to overcome by trained annotators.

For the posterior 746 sections, ROIs were not annotated prior to imaging. Instead, taking advantage of consistent section placement, we created a “blind” ROI that was ∼1.75× the width of the previously placed ROIs, which added imaging time but captured all the tissue. This approach yielded the same results as placing smaller ROIs prior to imaging.

### Image data alignment

Raw images were aligned using a custom pipeline based on AlignTK (https://mmbios.pitt.edu/aligntk-home) on the O2 computing cluster at Harvard Medical School as described previously (Bock et al., 2011; Nguyen et al., 2023; Phelps et al., 2021; Tobin et al., 2017). First, camera image tiles were stitched into continuous two-dimensional montages, and then consecutive montages were aligned into a three-dimensional volume. Every 50th section was used as a global constraint on the full dataset’s alignment. For sections with small artifacts that would warp or misalign with the standard elastic alignment, rigid constraints were applied (absolute_maps in AlignTK), and manually placed control points were used to map corresponding features in neighboring sections.

### Ribbon synapse manual imaging at 25,000× magnification

We performed higher-resolution re-imaging to examine the ultrastructure of ribbon synapses in greater detail. We targeted 18 out of 57 ribbons, which appeared to exhibit vesicle docking to the presynaptic membrane. Following 25 months in a dessicator (Secador, EMS), the reel was reloaded into the microscope. Single camera images at 25,000× were captured manually. Background was subtracted (rolling ball radius 50 pixels, light background) and local contrast was enhanced (histogram equalization, saturated pixels 0.4%) in ImageJ (version 1.52a) at a resolution of 0.4 × 0.4 nm per pixel.

### Anatomical nomenclature

Throughout this work, we use the nomenclature proposed for glomeruli in (Herre et al., 2022). We refer to the maxillary palp glomeruli as Glomerulus 1, 2, and 3. These glomeruli are referred to in previous literature as medio-dorsal (MD) glomeruli MD1, MD2, and MD3, respectively (Ignell et al., 2005; Shankar and McMeniman, 2020). The glomeruli in *Ae. aegypti* were originally named (Ignell et al., 2005) using a set of coordinate axes that is different from what is now commonly used in *D. melanogaster* (Ito et al., 2014), and these glomeruli are not notably dorsal in the coordinate space that we use here. We have therefore chosen the naming scheme that refers to them as Glomerulus 1, 2, and 3 (see (Herre et al., 2022) for a more complete description of antennal lobe glomeruli nomenclature).

### Reconstruction of Ae. aegypti neurons

Neurons were manually reconstructed in the EM dataset as described previously (Lee et al., 2016; Phelps et al., 2021; Tobin et al., 2017). We imported the data into Collaborative Annotation Toolkit for Massive Amounts of Image Data (CATMAID, release 2018.11.09) (Saalfeld et al., 2009) for distributed web-based reconstruction and annotation. To reconstruct neurons, we manually placed points, or nodes, down the center of each processes, creating skeletonized models of neuron morphology.

We identified the maxillary palp nerve bundle on the right side of the brain, with corroboration from light-level images, then completed tracing of 45 Glomerulus 1 OSNs, 45 Glomerulus 2 OSNs, and 46 Glomerulus 3 OSNs with axons in the bundle. For each OSN, the ventroposterior point of entry into the dataset was tagged as the root node, and the first branch point of the axonal arbor was tagged “first branch point”. The OSNs’ first branch point was typically close to where the OSN entered and started to branch and ramify in its home glomerulus. We identified the uPN axons in the projection nerve bundle (iACT, **Supplementary** Figure 1B) from the antennal lobe to higher brain centers. Tracing these back until they entered glomeruli, we found one densely branching uPN innervated each of the three maxillary palp glomeruli. PNs projecting through other tracts were not included in this analysis. Each uPN had a large-caliber process leading laterally to a cortical soma cluster, though only Glomerulus 1 uPN’s soma was included in the bounds of the dataset. An mPN was identified by an axon in the iACT, a process projecting laterally towards a soma, and innervation of two different antennal lobe glomeruli.

In OSNs and PNs, T-bar synapses were identified and annotated using ultrastructural criteria: a presynaptic T-bar, presynaptic vesicles, synaptic cleft, and postsynaptic densities (Prokop and Meinertzhagen, 2006; Wagh et al., 2006). We identified *en face* cut synapses when a section contained a presynaptic specialization and an adjacent section(s) contained vesicle pools, and putative postsynaptic cells contained postsynaptic densities. Putative ribbon synapses consisted of a presynaptic density oriented non-parallel to the presynaptic membrane that was surrounded on all sides by vesicles (**Supplementary** Figure 10). The vast majority occurred at lower branch orders (**Figure 6, Supplementary** Figures 10-11). Ribbon synapses were differentiated from obliquely cut synapses by vesicle pools present on the same sections as the presynaptic density and the lack of putative postsynaptic cells on sections adjacent to the presynaptic density.

To increase our annotation accuracy, each fully reconstructed OSN and uPN was proofread by a minimum of three trained annotators, who marked all synapses with a connector node and all postsynaptic partners with a new skeleton. We reconstructed all postsynaptic partners of the thirty fully reconstructed OSNs, ten randomly selected for each glomerulus, to identify their cell type. To empirically validate that the selected OSNs morphologies were representative, we used NBLAST to generate similarity scores for all OSNs in each glomerulus (Costa et al., 2016). LNs, or multiglomerular cells, were defined as reconstructed cells that extended beyond the home glomerulus. Some presynaptic specializations (<10%) appeared to contact glia or to interstitial space between cells.

### Connectivity analysis

Analysis was performed in Python using pymaid (version 2.4.0, https://pymaid.readthedocs.io) and NAVis (version 1.5.0, https://navis.readthedocs.io). Data visualization and plotting were performed using matplotlib (version 3.8.3) and seaborn (version 0.13.2) packages and statistics were performed using SciPy (version 1.12.0), with posthoc tests performed using scikit-posthocs (version 0.9.0).

To measure cable length and distances between synapses and branch points, we computed geodesic or “along the arbor” distances. Because the opportunities for neurons to be synaptically connected is limited to regions where neuron processes are close enough to make synapses, we control for this by computing synapse density. We define synapse density as the number of presynaptic output synapses normalized by the cable overlap, which is the length of presynaptic neurite within sufficiently close proximity (< 2 µm) of the postsynaptic neurite to make a synapse.

To control for differences in neurite lengths from each glomerulus to where axons reach the boundaries of the EM volume, we used two approaches to restrict our analysis to within glomeruli themselves. First, we generated meshes that defined the 3D boundary of the glomeruli and limited our analyses to neurites and synapses within the bounds of the 3D meshes. Second, for OSNs, we restricted our analysis to branches and synapses distal from the first branch point of the OSN axon, which was typically near where the axon enters the glomerulus.

### Axogram and dendrograms

Axograms and dendrograms were generated with Neuroboom (Felsenberg et al., 2018) (version 0.3.55, https://github.com/markuspleijzier/neuroboom). OSNs were plotted vertically (dot progression), whereas PNs were plotted radially (neato progression) to embed their more complex arbors.

### Additional technical notes on reconstructed neurons

We identified a single putative mPN that sparsely innervates Glomerulus 1 and another unidentified glomerulus. This neuron has an axon tract that exits the antennal lobe in iACT, and may be an mPN. The branches were reconstructed, but it was not used in inter-glomerular comparative analyses because we were unable to do a complete survey of mPNs without a larger EM volume. We do not rule out the existence of other PNs that may exit through separate tracts (Bates et al., 2020; Liang et al., 2013; Tanaka et al., 2012).

### Analysis of D. melanogaster neurons in Flywire

To compare *Ae aegypti* glomerular wiring and connectivity to that of *D. melanogaster*, we first identified antennal lobe glomeruli in the Flywire dataset (Dorkenwald et al., 2024, 2022). We then proofread OSNs of the CO2-selective glomerulus V and appetitive glomerulus DM1 on the right side of the brain for merge and split errors; we selected the right side was because at the time it contained better-proofread neurons. Subsequent proofreading by the FlyWire consortium completed the rest of the OSNs and PNs (Dorkenwald et al., 2024; Schlegel et al., 2024). We analyzed fly OSNs, PNs, and their connectivity in the right hemisphere using the glomerular boundary meshes made previously (Schlegel et al., 2021). We used fafbseg (version 3.0.10; https://fafbseg-py.readthedocs.io) to pull data from the 3D reconstructions and the skeletonization function to generate skeletons from 3D neuron meshes. Specifically, OSNs were identified using the hemibrain_type field for all glomeruli except VC3, VC5, and VM6 which utilized the cell_type field due to changes in FlyWire’s data structure. Additionally, we removed cells (N=33) labeled as OSNs from the analysis if they did not have any synapses onto a PN. For the purposes of comparison, we used a previously compiled list of uPNs (Bates et al., 2020). We then used the flywire.skid_to_id function to recover the root_id in the FlyWire dataset.

In Flywire, a synapse prediction network (Buhmann et al., 2021) assigns each predicted synapse a cleft score, higher cleft scores reduce the chance of a false positive result, though the likelihood of a false negative result also increases. We used a cleft score of 65 for the purposes of this analysis. To remove instances of nonexistent autapses we set the parameter filter=True regardless of cleft score.

### Quantification and statistical analysis

Unless otherwise stated, we used two-tailed Kruskal-Wallis tests and Dunn’s posthoc tests with Bonferroni correction for pairwise comparisons. We implemented an algorithm (Piepho, 2004) that utilizes pairwise comparisons to identify significance groupings (**Supplemental Figures 7-8**). We use the non-parametric Kruskal-Wallis test to statistically compare across more than two groups because we do not assume normality and it is more robust to outliers. Some *D. melanogaster* glomeruli exhibit extreme values (**Figures 3-4 and Supplemental Figures 7-8**). Since normality was not assumed, a formal statistical test for outliers was not applicable. However, we inspected all data for points where synapse density was greater than two median absolute deviations (2×MADs) from the median. All feedforward synapses that were 2xMAD from the median that we visually inspected were verified as synapses and these data were included in all analyses. We identified only 4 false-positive recurrent synapses among the outlier population. To most accurately reflect the morphology these synapses were removed from the data presented throughout the paper. This difference did not change any of the statistically significant findings The locations of the synapses in the FlyWire dataset that were removed are:

### Network model

We simulated our model in Python using custom software. A notebook for simulating the modeling results can be found at: https://github.com/htem/aedes_public. Model parameters were selected based on previous work (Hallem and Carlson, 2006; Luo et al., 2010; Olsen et al., 2010). We used values of σ=12 spikes/s and *Rmax* =165 spikes/s based on OSN and PN models fit to experimental data (Olsen et al., 2010). The exponent 1.5 was used to simulate CO2 in the presence of other odors (‘background odors’) that broadly activated many glomeruli (Olsen et al., 2010). We set *s* = 50 spikes/s. This value was consistent with models (Luo et al., 2010; Olsen et al., 2010), and also with a weighted average OSN response of 43 sp/s calculated from 186 odor responses from 24 *D. melanogaster* glomeruli (Hallem and Carlson, 2006). To select PN and OSN threshold values for our detection task, we used previous experimental findings. To estimate the PN threshold, we computed the average PN response across a distribution of odors (Hallem and Carlson, 2006) and selected this threshold to be two standard deviations larger than this mean. We assumed PN responses around this mean were Gaussian distributed with previously derived variance parameters (Luo et al., 2010). To compute the OSN threshold of 35 sp/s, we considered experiments that found that in an environment devoid of background CO2, OSNs in *Ae. aegypti* were unresponsive to CO2 concentrations less than 200 ppm CO2 but responded at a rate of approximately 35 sp/s for concentrations of 300 ppm, which they noted corresponds to the ambient CO2 level in natural environments (Grant et al., 1995). Additional, experimental work also found OSN response rates of around 35 sp/s for CO2 concentrations 200 ppm greater than background CO2 concentrations, for a variety of background levels (Majeed et al., 2014). Values of *g* identified in Figure 6D were found using scipy’s root finding method, root_scalar.

## Acknowledgements

We thank Leslie Vosshall and the Vosshall Lab at Rockefeller University for specimens and for initial support of the project; Mingguan Liu, Marine Nimblette, Elaina Phalen, Laurel Guo, Manuela Eroles, Karenna Ng, Genevieve Hulshof, Mark Larson, Katie Molloy, Nicholas Byrne, Olivia Sato, Brian Reicher, Catrin Zharyy, and Shuhan Xie for neuron reconstructions and proofreading; Steve Muscari for help with EM imaging; Kenneth Hayworth for help with X-ray microCT processing; Zhongyan Gong for help with EM alignment; Jasper Phelps for advice and EM dataset alignment; Aaron Kuan, Steve Muscari, Jeff Rhoades, Alex Bates, Philip Schlegel, Maria Ericsson, Elio Raviola, Pascal Kaeser, and Markus Pleizer for discussions, advice, and assistance; Brett Graham, Steve Muscari, Logan Thomas, and Kris Kim for hardware development and maintenance; and Ian Davison, Tyler Hill, Brittany Ahn, Garima Kohli and Lee lab members for feedback on the manuscript. We thank Larry Abbott for helpful comments and insights on the network model.

## Author contributions

J.B., W.A. D.G.C.H., M.A.Y. and W.C.A.L. conceptualized the project and designed experiments. D.G.C.H., M.A.Y. and W.C.A.L. prepared the EM specimens. D.G.C.H. performed GridTape sectioning. J.B. and W.C.A.L. performed EM imaging. J.B. performed and managed neuron tracing and data annotation. G.L., A.K., L.S.C., L.W., and S. P. performed neuron tracing. W.A., Y.A., G.L., and J.B. developed code for data analysis. W.A., J.B., and W.CA.L. performed data analysis. L.W. and L.S.C. generated illustrations. J.B., W.A., M.A.Y., and W.C.A.L. wrote the paper. **Competing interests:** W.C.A.L. and D.G.C.H. declare the following competing interest: Harvard University filed a patent application regarding GridTape (WO2017184621A1) on behalf of the inventors including W.C.A.L, D.G.C.H. and negotiated licensing agreements with interested partners. All other authors declare no competing interests.

## Data and materials availability

The aligned EM dataset is available on the web-based CATMAID platform and custom code is available at https://github.com/htem/aedes_public

---

**Supplementary Figure 1:**
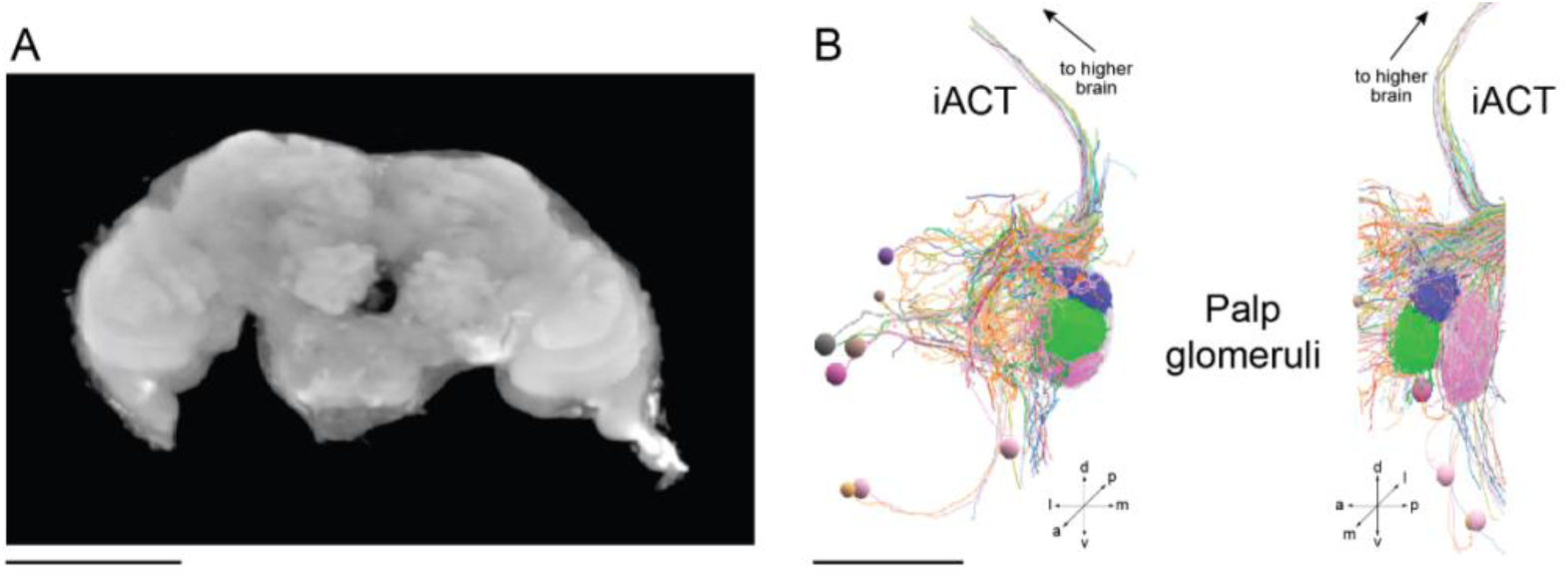
MicroCT and PNs with axons in iACT. (A) X-ray micro-computed tomography (microCT) of the *Ae. aegypti* brain analyzed in this study (anterior view). Scale bar 100 µm. (B) Coronal *(left)* and sagittal *(right)* views of all PNs on the left side of the brain that contained axons that fasciculate into the ipsilateral iACT. Ispilateral iACT uPNs were traced to completion while putative multiglomerular cells were not. a, anterior; d, dorsal; l, lateral; m, medial; p, posterior; v, ventral. Scale bar 50µm.

**Supplementary Figure 2:**
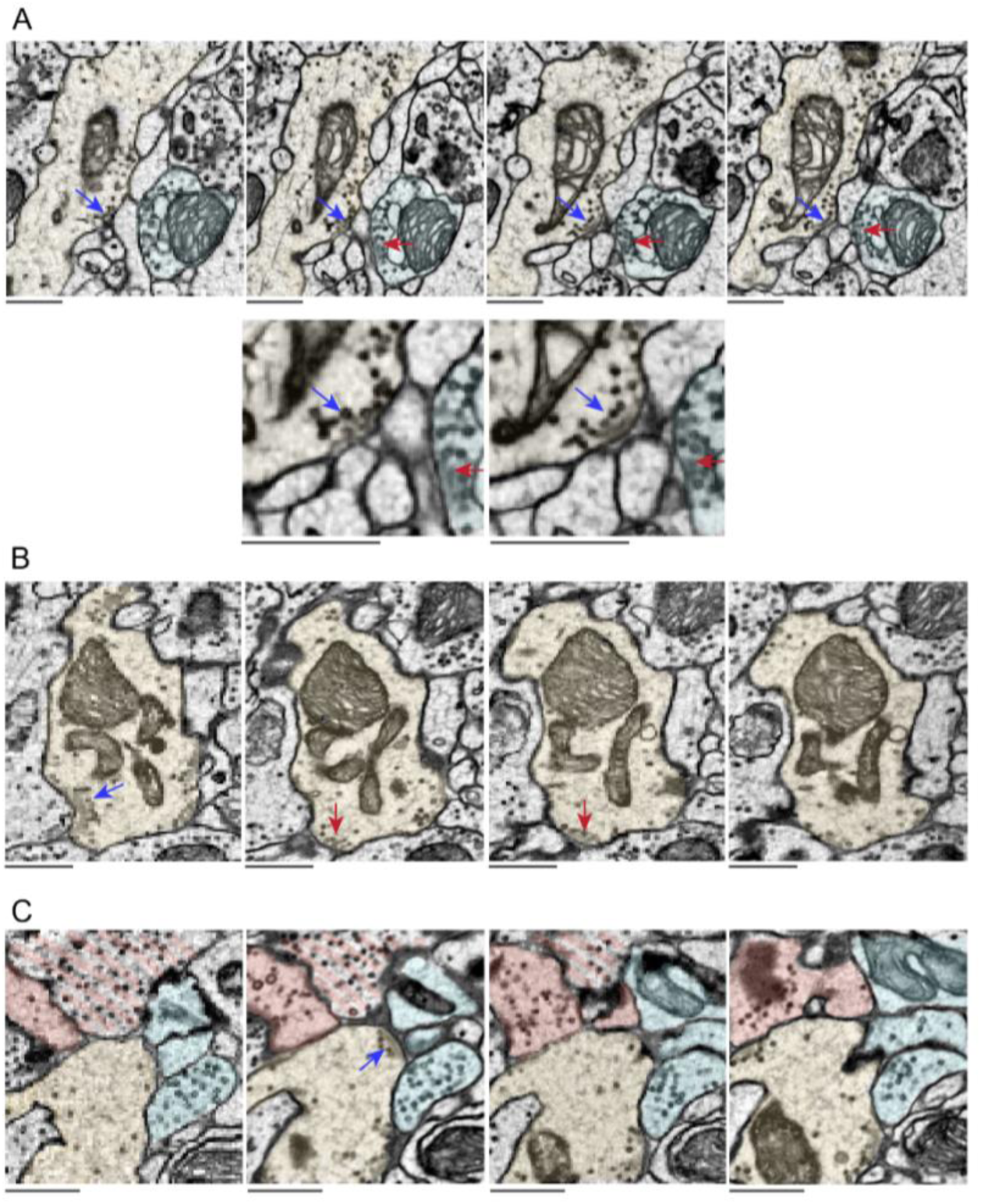
Examples of T-bar synapses and postsynaptic partners in *Aedes aegypti*. (A) Zoomed out (top) and zoomed in (bottom) canonical T-bar synapses, indicated by blue and red arrows respectively. Here, both synapses are characterized by the clustering of vesicles, postsynaptic densities, and presynaptic T-bar structures. (B) Example of one presynaptic cell making two closely located synapses. These are considered separately as one (blue arrow) synapse’s characteristics disappear for at least 40 nm before appearing for the subsequent nearby synapse (red arrow). (C) Example of one T-bar synapse (blue arrow) and its postsynaptic partners. Presynaptic cell in yellow, postsynaptic partners in blue. Partners not considered postsynaptic for this synapse - based on contact with the synaptic cleft - are marked in red. Scale bars 500 nm.

**Supplementary Figure 3:**
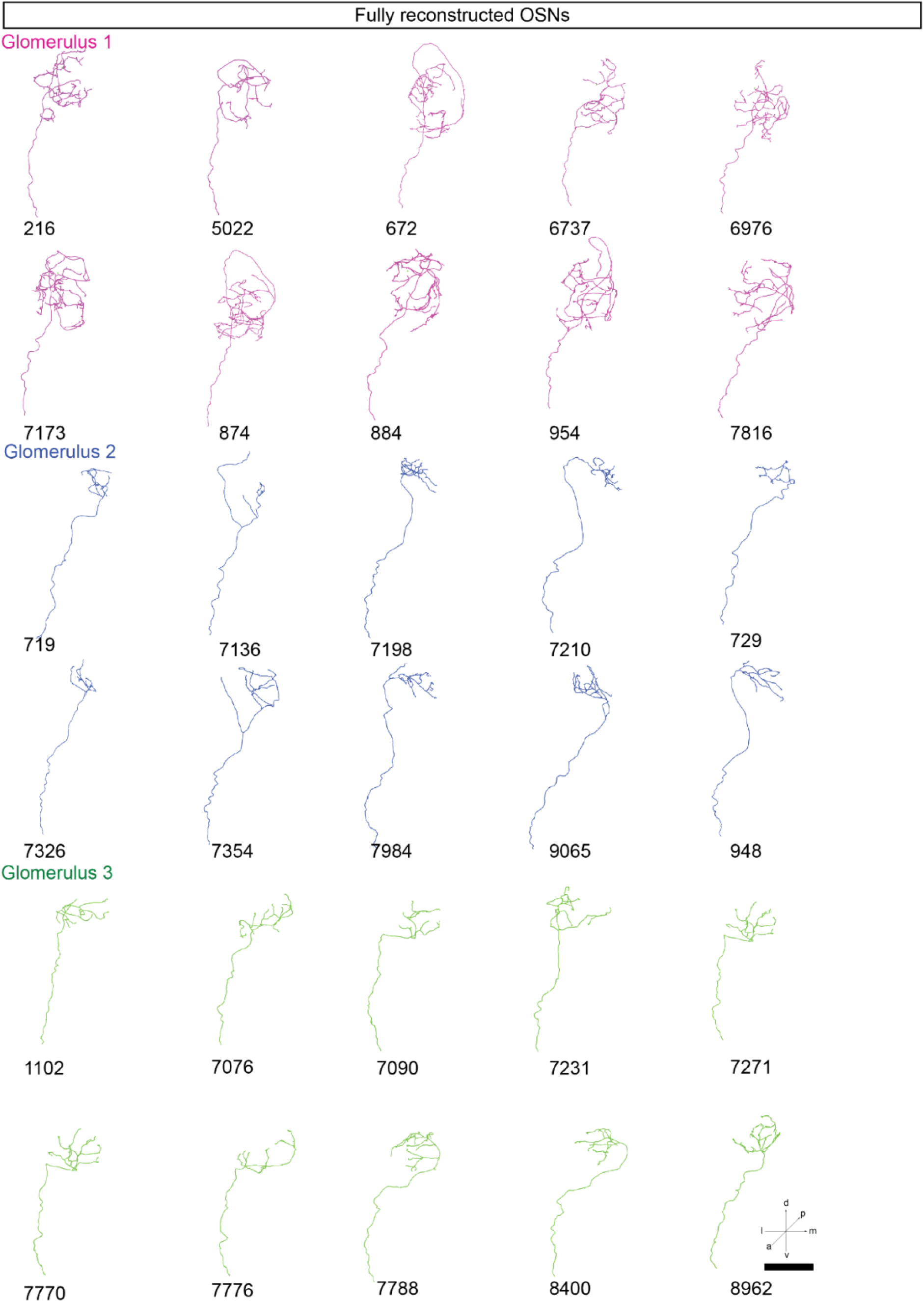
Reconstructed OSN morphologies. Skeletonized neuron morphologies are shown for the 10 OSNs fully reconstructed from Glomerulus 1, 2, and 3. Labels are neuron IDs corresponding to the CATMAID database. Neuron 7136 had a proximal branch that projects dorsoposteriorly toward higher brain areas. a, anterior; d, dorsal; l, lateral; m, medial; p, posterior; v, ventral. Scale bar 25 µm.

**Supplementary Figure 4:**
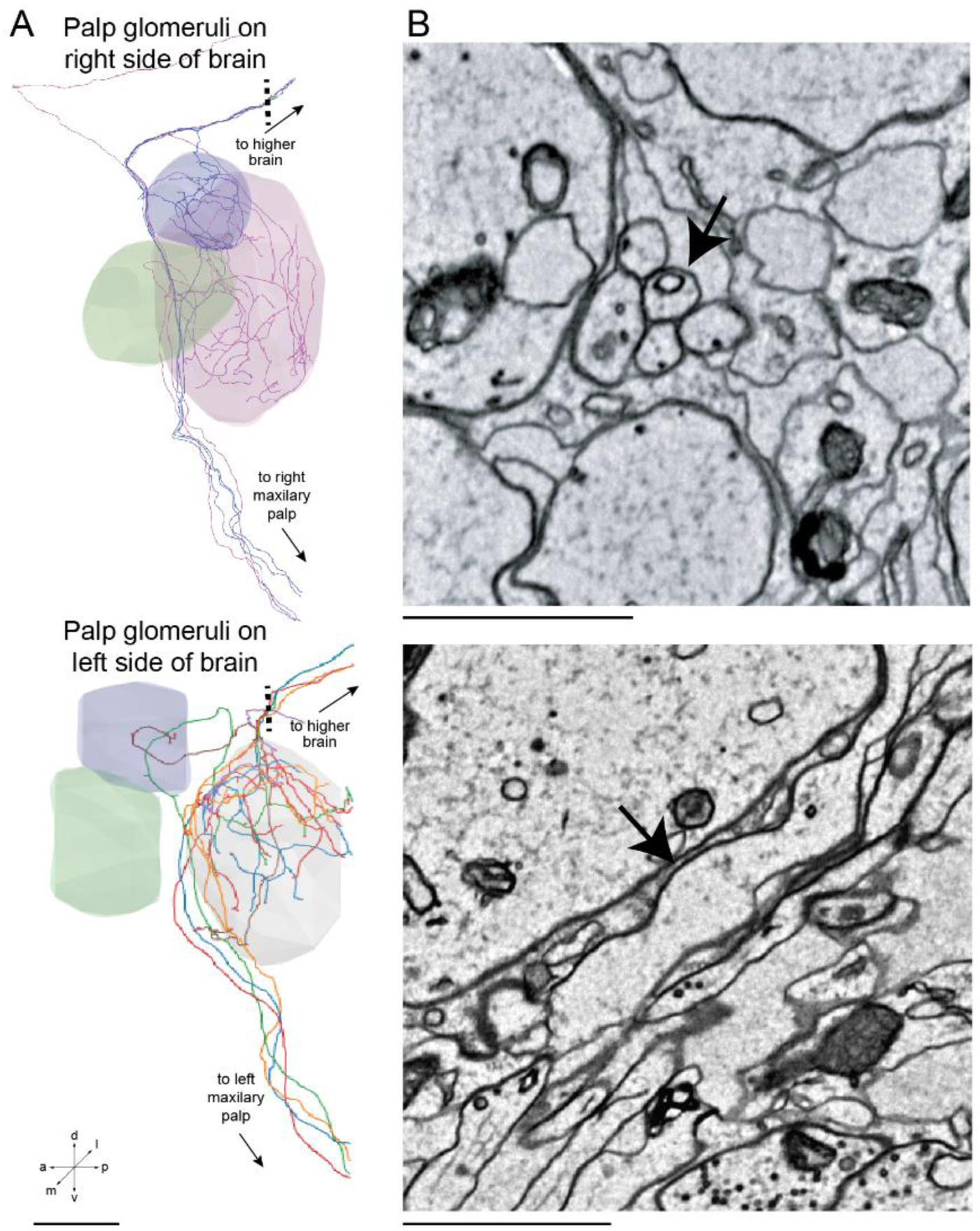
Dorso-posteriorly projecting tract of OSN processes co- fasciculates with the iACT. (A) Rendering of 5 OSNs (colored traces) innervating Glomerulus 1 and Glomerulus 2 (gray volumes represent glomerular boundaries; Glomerulus 3 is also depicted, but none of the extra- projectional OSNs innervate it) that send small-caliber neurites along the iACT (medial uPN tract, Supplementary Fig. 1B) to higher-order brain regions. *(top)* Right side of brain. *(bottom)* Left side of brain. Dashed lines indicate locations of EM micrographs shown in (B). a, anterior; d, dorsal; l, lateral; m, medial; p, posterior; v, ventral. Scale bar 25 µm. (B) EM micrograph at dashed line locations in (A) Co-fasciculating neurites indicated by arrows. Scale bars 1 µm.

**Supplementary Figure 5:**
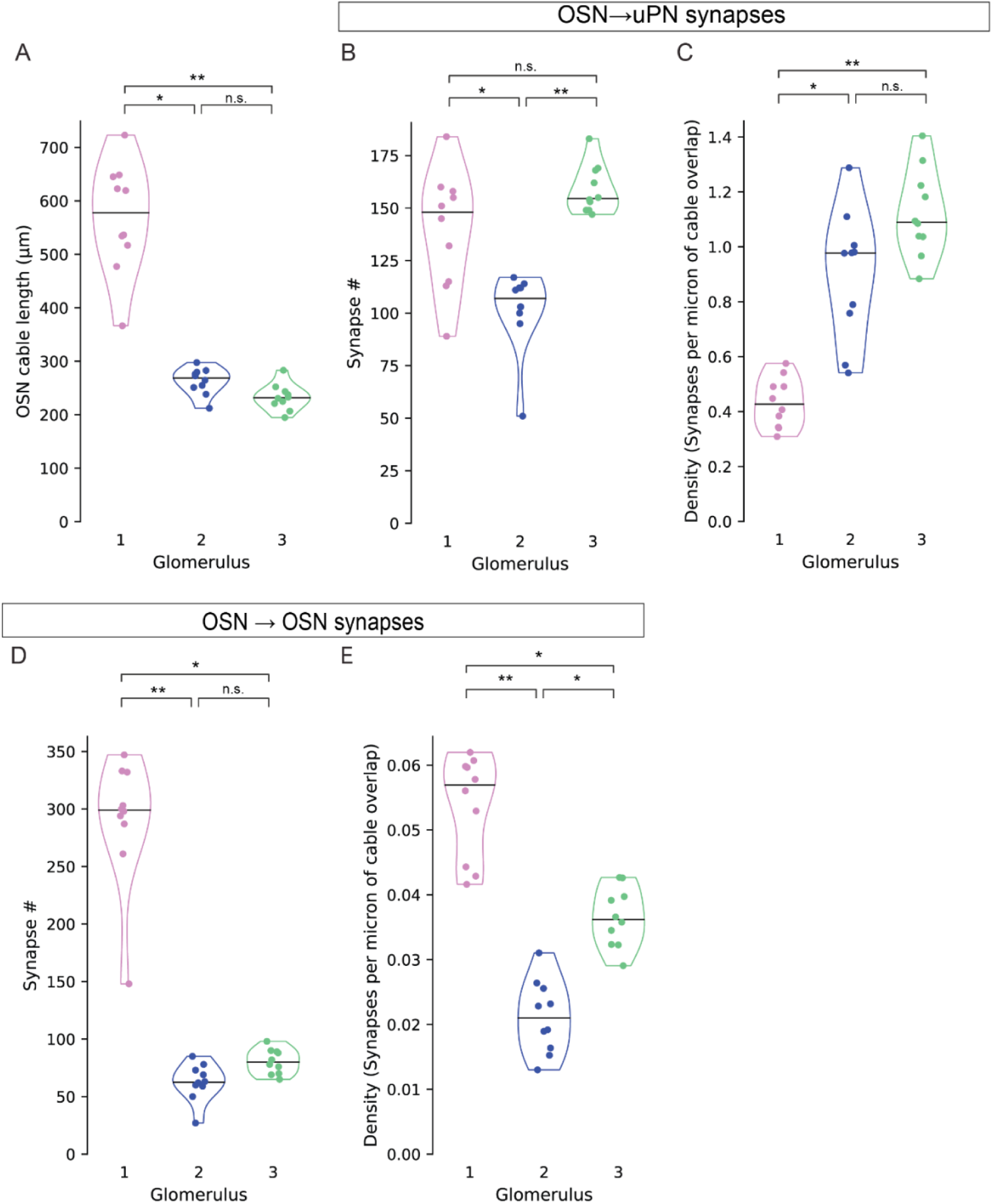
Analyses of connectivity including extraglomerular OSN and uPN axons. (A) OSN cable length within each glomerulus. (Kruskal-Wallis test (two-tailed), p = 1.81 x 10^-5^, pairwise comparisons using Dunn’s post-hoc test with Bonferroni correction: n.s., p > 0.05; * p ≤ 0.01; ** p ≤ 0.001). (B) Number of OSN-to-uPN synapses (Kruskal-Wallis test (two-tailed), p = 0.0002, pairwise comparisons using Dunn’s posthoc test with Bonferroni correction: n.s., p > 0.05; * p ≤ 0.01). (C) Density of OSN-to-uPN synapses (Kruskal-Wallis test (two-tailed), p = 3.61 x 10^-5^, pairwise comparisons using Dunn’s post-hoc test with Bonferroni correction: n.s., p > 0.05; * p ≤ 0.05; ** p ≤ 0.001). (D) Total number of outgoing OSN-to-OSN synapses restricted to within glomerular boundaries (Kruskal-Wallis test (two-tailed), p = 1.28 x 10^-5^, pairwise comparisons using Dunn’s post-hoc test with Bonferroni correction: n.s., p > 0.05; * p ≤ 0.05; ** p ≤ 0.001). (E) Density of OSN-to-OSN synapses to within glomerular boundaries (Kruskal-Wallis test (two-tailed), p = 3.65 x 10^-6^, pairwise comparisons using Dunn’s post-hoc test with Bonferroni correction: n.s., p > 0.05; * p ≤ 0.01).

**Supplementary Figure 6:**
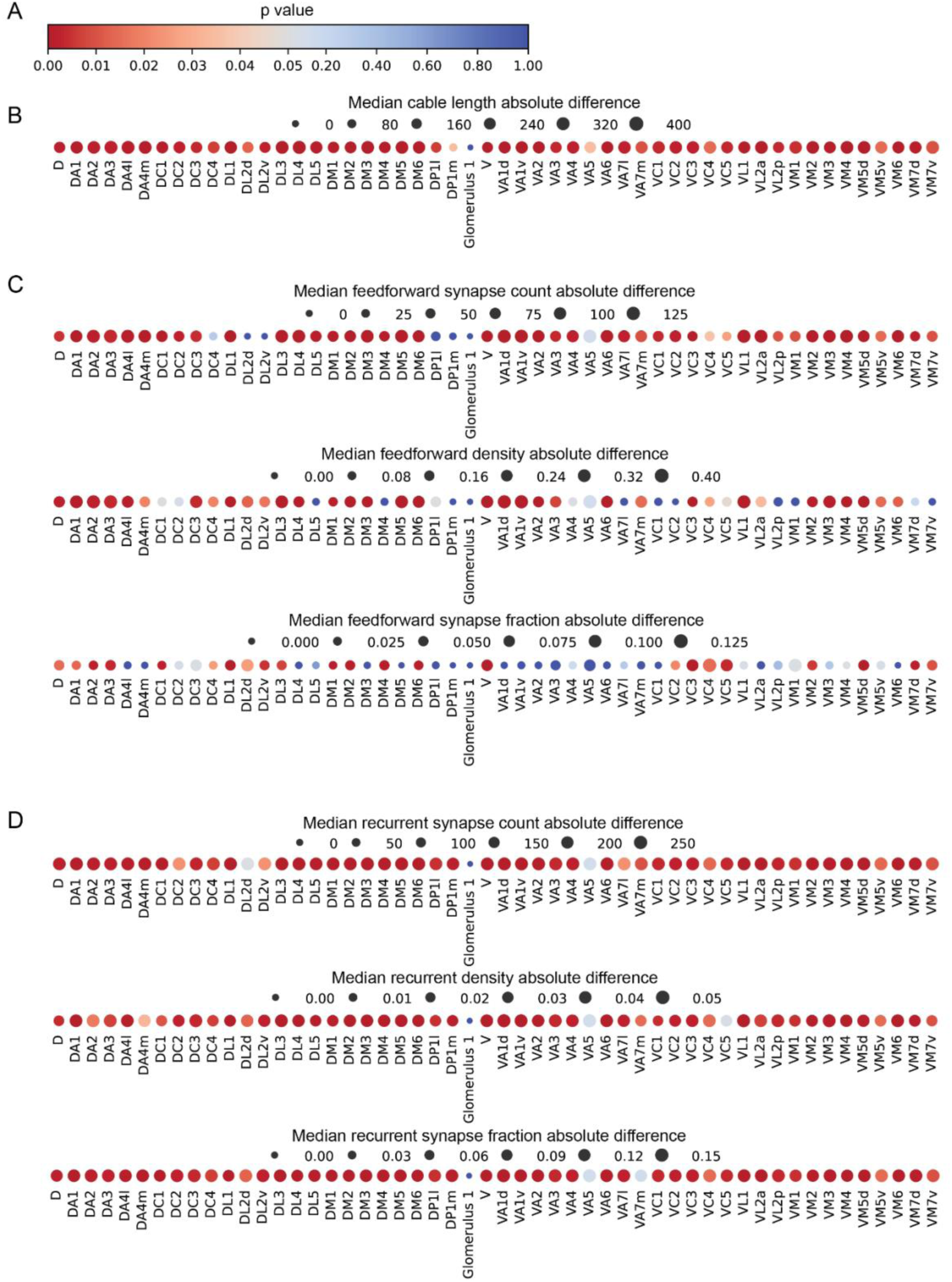
Visualization of statistical significance and median difference of morphological metrics of *D. melanogaster* and Glomerulus 1. Representation of each glomerulus is composed of two parts: size and color. The size of each point is the linearly scaled absolute difference of each metric. The color of each point is the significance of the pairwise statistical test for each glomerulus. (A) Color bar where significance from 0 to 0.05 is scaled at a different linear rate than from 0.05 to 1 to emphasize different degrees of significance. Red hues represent significance and blue hues represent non-significance.(B) Median difference of cable length within each glomerulus from Glomerulus 1, significance as indicated in (A). (C) Median difference of feedforward metrics of each glomerulus from Glomerulus 1, significance as indicated in (A). (D) Median difference of recurrent metrics of each glomerulus from Glomerulus 1, significance as indicated in (A).

**Supplementary Figure 7:**
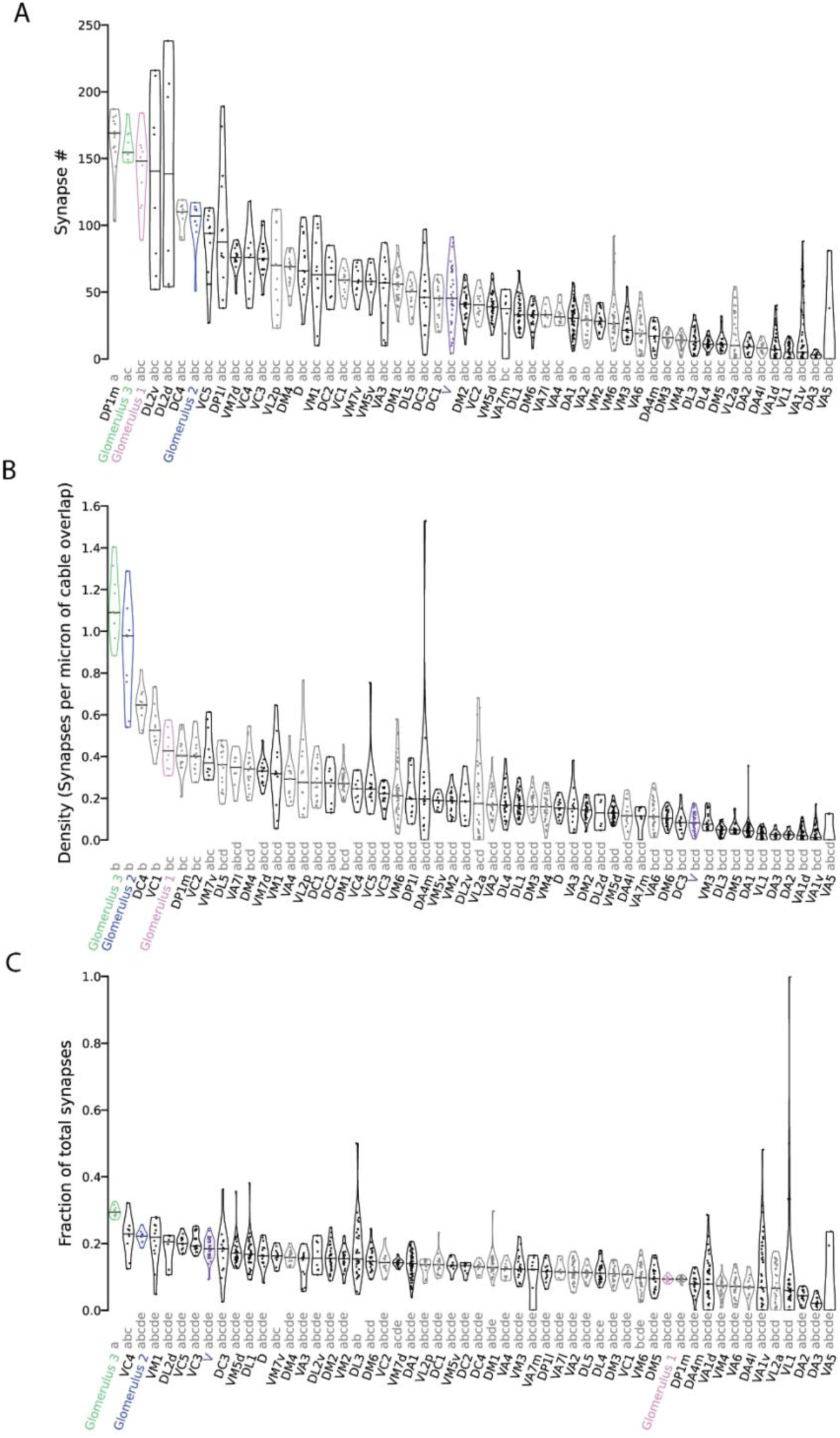
Pairwise analysis of feedforward connectivity for all glomeruli. (A) Number of OSN-to-uPN synapses restricted to within glomerular boundaries (Kruskal-Wallis test (two-tailed), p = 9.21 x 10^-140^, different letters mark whether glomeruli are significantly different using Dunn’s posthoc test with Bonferroni correction: p ≤ 0.01). (B) Density of OSN-to-uPN synapses within glomerular boundaries (Kruskal-Wallis test (two-tailed), p = 1.32 x 10^-137^, compact letter display as in (A)). (C) OSN-to-uPN synapses restricted to within glomerular boundaries as a fraction of overall OSN output synapses (Kruskal-Wallis test (two-tailed), p = 1.63 x 10^-91^, compact letter display as in (a)).

**Supplementary Figure 8:**
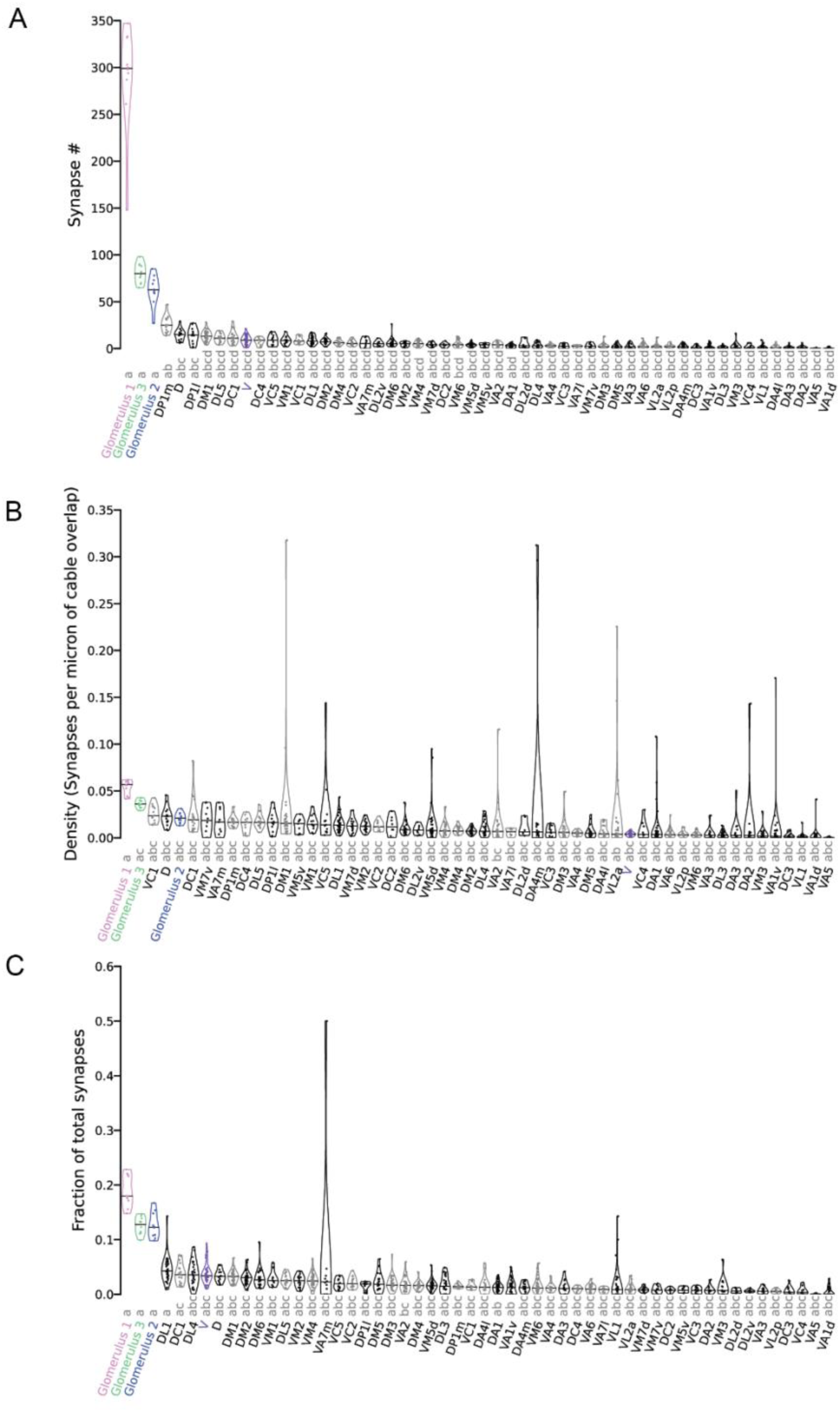
Analyses of recurrent connectivity for all glomeruli. (A) Number of OSN-to-OSN synapses restricted to within glomerular boundaries (Kruskal-Wallis test (two-tailed), p = 7.88 x 10^-105^, different letters mark whether glomeruli are significantly different using Dunn’s posthoc test with Bonferroni correction: p ≤ 0.01). (B) Density of OSN-to-OSN synapses within glomerular boundaries (Kruskal-Wallis test (two-tailed), p = 6.96 x 10^-78^, compact letter display as in (A)). (C) OSN-to-OSN synapses restricted to within glomerular boundaries as a fraction of overall OSN output synapses (Kruskal-Wallis test (two-tailed), p = 1.03 x 10^-79^, compact letter display as in (a)).

**Supplementary Figure 9:**
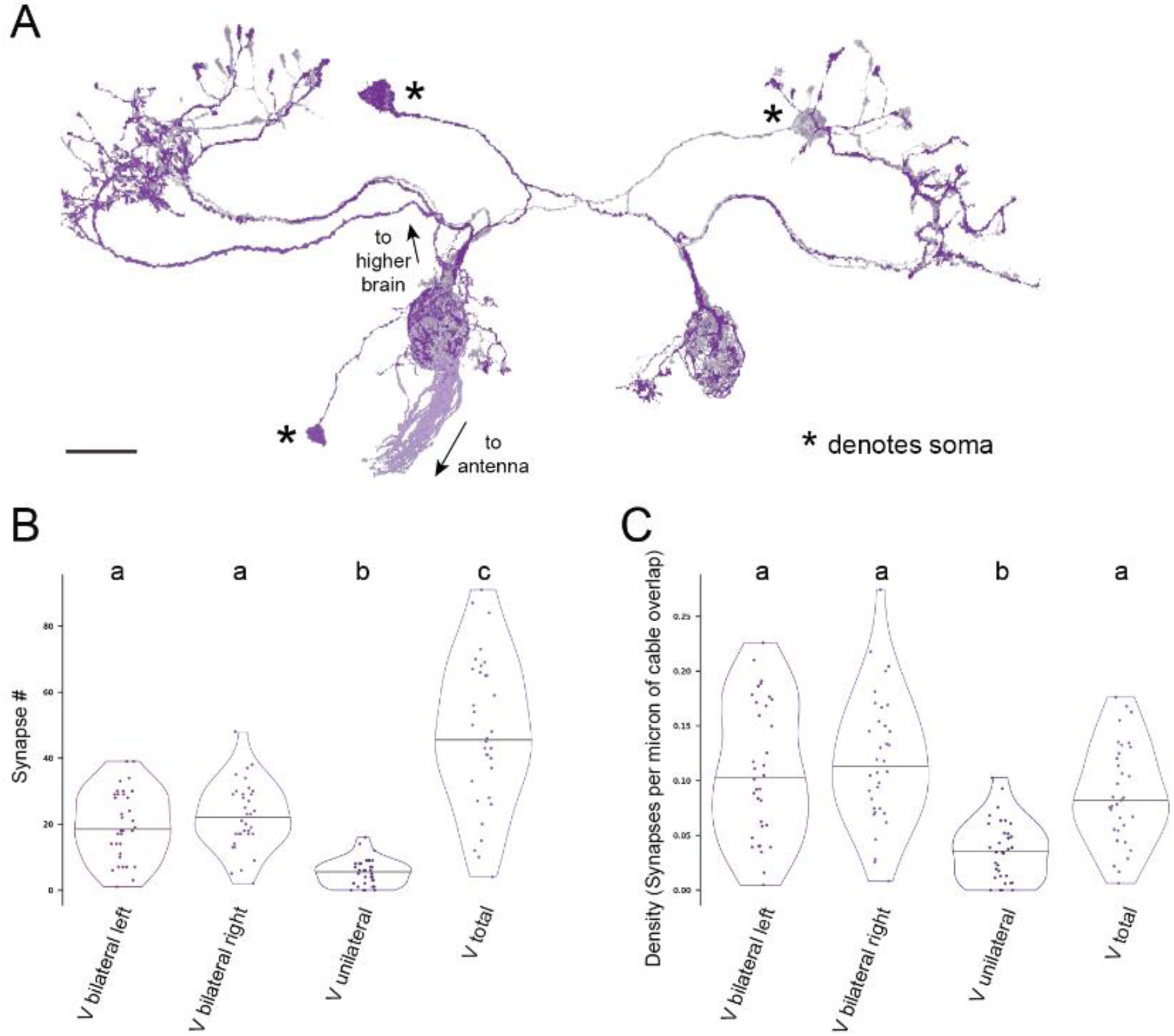
Reconstructions of OSNs and uPNs of *D. melanogaster* glomerulus V. (A) V OSNs and uPNs reconstructed in Flywire (Bates et al., 2020; Dorkenwald et al., 2024; Schlegel et al., 2024). Lighter shades of orange and purple correspond to OSNs, while darker shades correspond to uPNs. Scale bars 50 µm. (B) Number of OSN to uPN synapses restricted to within glomerular boundaries (Kruskal-Wallis test (two-tailed), p = 5.17 x 10^-17^, different letters mark whether glomeruli are significantly different using Dunn’s posthoc test with Bonferroni correction: p ≤ 0.01). (C) Density of OSN to uPN synapses within glomerular boundaries (Kruskal-Wallis test (two-tailed), p = 1.62 x 10^-09^, letters display significant differences as in (B)).

**Supplementary Figure 10:**
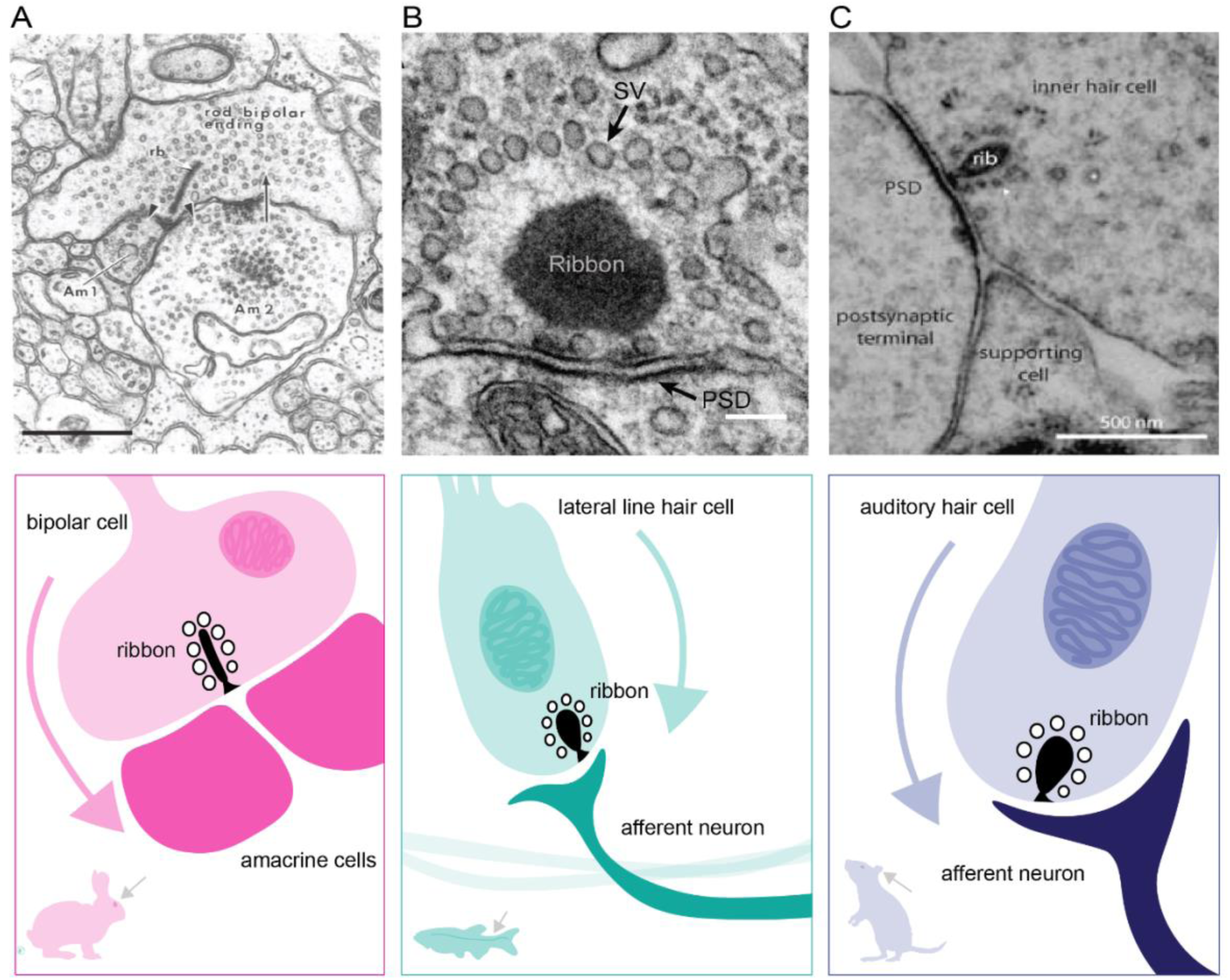
Examples of ribbon synapses in vertebrates (A-C) Examples EM micrographs *(top)* and schematics *(bottom)* of ribbon synapses across vertebrate species. (A) Ribbon (rb) synapse (arrowheads) in rabbit retina between rod bipolar cells and amacrine (Am) cells (Raviola and Dacheux, 1987). Arrow shows reciprocal synapse from amacrine cell back to the bipolar ending. Scale bar 200 nm. (B) Ribbon synapse in a zebrafish lateral line hair cell (Kindt and Sheets, 2018). SV, synaptic vesicle; PSD, postsynaptic density. Scale bar 100 nm. (C) Ribbon (rib) synapse in mouse auditory hair cell (Nouvian et al., 2006). Scale bar 500 nm.

**Supplementary Figure 11.**
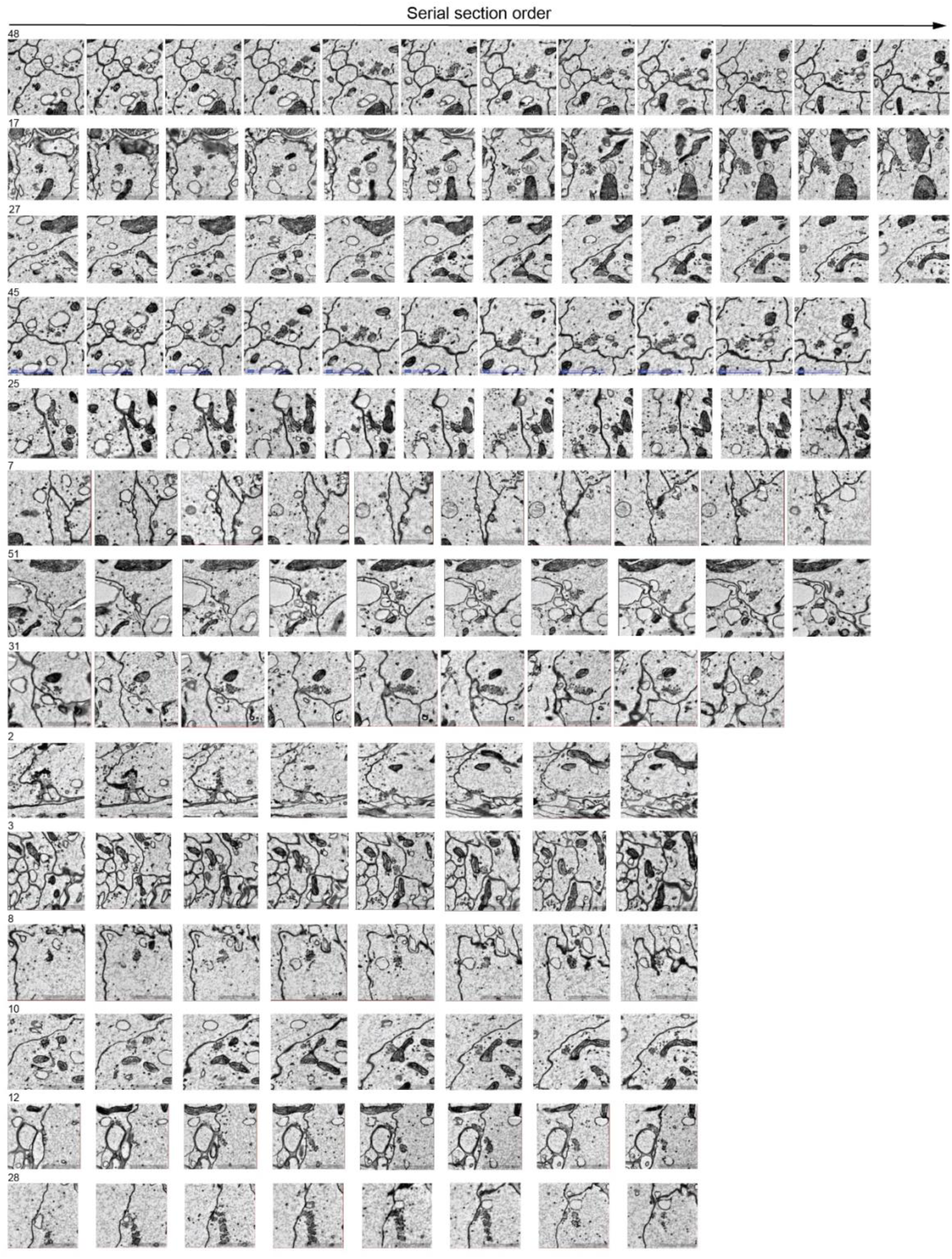
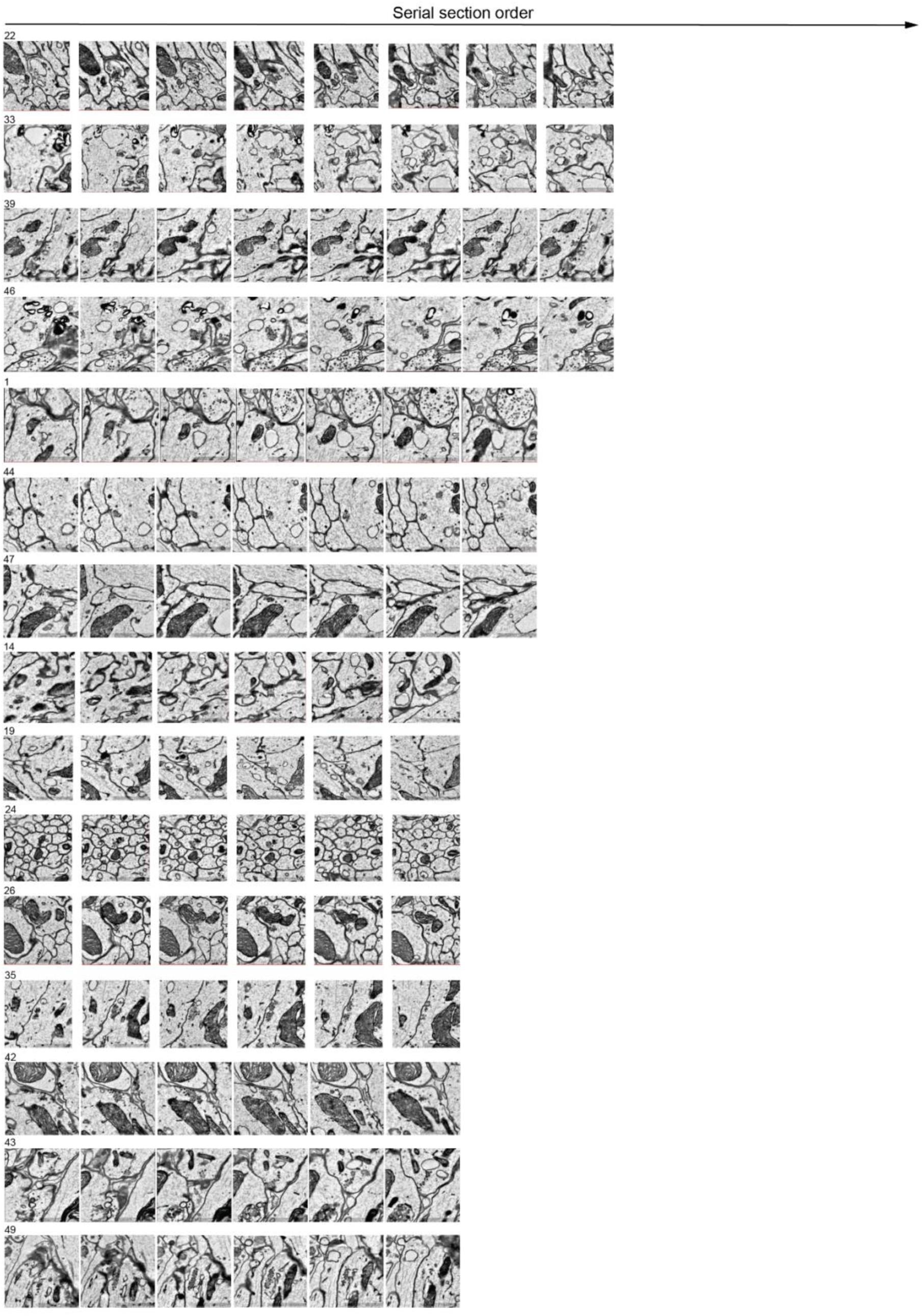
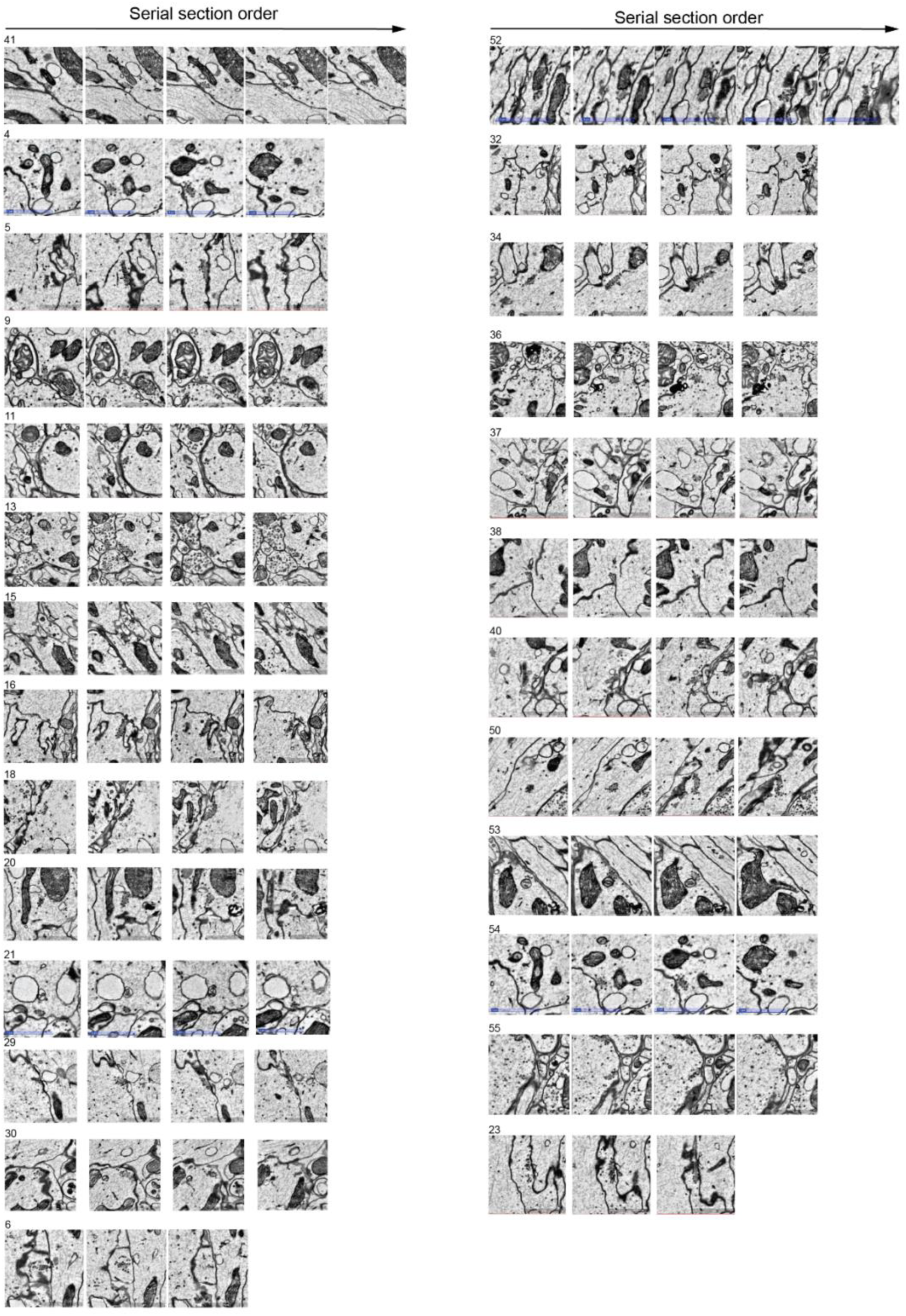
(3 pages): Serial-sections of ribbon-like structures. Serial electron micrographs through ribbon-like structures in Glomerulus 1 OSNs. Labels are connector IDs corresponding to the CATMAID database. Scale bars 1 µm.

